# *SPHERIOUSLY?* The challenges of estimating spherical pore size non-invasively in the human brain from diffusion MRI

**DOI:** 10.1101/2020.11.06.371740

**Authors:** Maryam Afzali, Markus Nilsson, Marco Palombo, Derek K Jones

## Abstract

The Soma and Neurite Density Imaging (SANDI) three-compartment model was recently proposed to disentangle cylindrical and spherical geometries, attributed to neurite and soma compartments, respectively, in brain tissue. The approach could also enable estimation of microstructure parameters such as the apparent size (radius) of the soma. There are some recent advances in diffusion-weighted MRI signal encoding and analysis (including the use of multiple so-called ‘b-tensor’ encodings and analysing the signal in the frequency-domain) that have not yet been applied in the context of SANDI. In this work, using: (i) ultra-strong gradients; (ii) a combination of linear, planar, and spherical b-tensor encodings; and (iii) analysing the signal in the frequency domain, three main challenges to robust estimation of soma size were identified:

First, the Rician noise floor in magnitude-reconstructed data biases estimates of soma properties in a non-uniform fashion. It may cause overestimation or underestimation of the soma size and density. This can be partly ameliorated by accounting for the noise floor in the estimation routine.

Second, even when using the strongest diffusion-encoding gradient strengths available for human MRI, there is an *empirical* lower bound on the spherical signal fraction and pore-size that can be detected and estimated robustly. For the experimental setup used here, the lower bound on the signal fraction was approximately 10%. We employed two different ways of establishing the lower bound for spherical radius estimates in white matter. The first, examining power-law relationships between the DW-signal and diffusion weighting in empirical data, yielded a lower bound of 7 *μm*, while the second, pure Monte Carlo simulations, yielded a lower limit of 3 *μm* and in this low radii domain, there is little differentiation in signal attenuation.

Third, if there is sensitivity to the transverse intra-cellular diffusivity in cylindrical structures, e.g., axons and cellular projections, then trying to disentangle two diffusion-time-dependencies using one experimental parameter (i.e., change in frequency-content of the encoding waveform) makes spherical pore-size estimates particularly challenging.

We conclude that due to the aforementioned challenges spherical pore size estimates may be biased when the corresponding signal fraction is low, which must be considered when using them as biomarkers in clinical/research studies.

## 1. Introduction

Diffusion magnetic resonance imaging (dMRI) is a non-invasive technique widely used to study brain microstructure *in vivo*. Most dMRI methods are based on the conventional Stejskal-Tanner experiment (Stejskal and Tanner, 1965) that applies a pair of pulsed field gradients along a single axis for each signal preparation, which we refer to here as ‘linear’ encoding. Using linear encoding, disentangling different microstructural properties such as their size, shape, and orientation is far from trivial (Lampinen et al., 2017; Novikov et al., 2019). Such features may be entangled in the encoding process resulting in low specificity in their estimation. This is particularly problematic in dMRI where the image voxel is on the scale of a millimeter, and can therefore contain multiple microenvironments.

Biophysical modeling is often used to tackle the inverse problem of inferring relevant tissue features (such as cell size, shape, and orientation) from the measured dMRI signal (Mitra et al., 1992; Wiegell et al., 2000; Stanisz et al., 1997; Zhang et al., 2012; Assaf et al., 2008). Most contemporary dMRI models for neural tissue share some common assumptions and features. First, they separate the tissue into intra- and extra-neurite compartments. Second, the exchange between the compartments is considered to be negligible, such that each compartment has a fixed and time-invariant signal fraction *f_i_*, where ∑_*i*_ *f_i_* = 1. Third, most models treat the intra-neurite compartment as a ‘stick’ - that is a compartment in which the diffusivity perpendicular to the long axis of the compartment is assumed to be effectively zero. This assumption is based on the lack of sensitivity to the neurite diameter (Nilsson et al., 2017). Different models have been used for the orientational dispersion of the compartments, represented by an orientation distribution function (ODF). Some models consider only purely parallel orientations (i.e., a delta function on the sphere ODF) (Stanisz et al., 1997; Assaf et al., 2008) while others use spherical harmonics (Jespersen et al., 2007) or a function such as the Watson distribution (Zhang et al., 2012) to characterise orientation dispersion. A Gaussian anisotropic representation is most often used for the extra-neurite compartment. Its orientation is determined by the mean of the fiber ODF and is characterized by axial and radial diffusivities. In white matter, the extra-neurite compartment largely corresponds to the extra-axonal space (and possibly extra-astrocytic-process space). However, in gray matter, the extra-neurite compartment is comprised of the extracellular space (where the Gaussian representation is likely valid) and an additional contribution from the water restricted inside isotropic cellular structures, such as the cell bodies (soma) (Palombo et al., 2020).

Recent studies have shown that the two-compartment model is not a good representation of the signal in gray matter (Afzali et al., 2020a,c; McKinnon et al., 2017; Veraart et al., 2019; Palombo et al., 2018a; Henriques et al., 2019; Jespersen et al., 2019). This can be due to non-negligible water exchange processes occurring between intra- and extra-cellular compartments and between different intracellular compartments (Veraart et al., 2018; Jelescu and Novikov, 2020), or the assumption that the water inside the soma behaves the same as water in extra-cellular space. (Palombo et al., 2018a,b). Addressing this model insufficiency, Palombo et al. (Palombo et al., 2020) first demonstrated with non-trivial numerical simulations that, under specific experimental conditions, the contribution of soma to the total intracellular dMRI signal can be disentangled from that of neurite, and then introduced a three-compartment model called Soma And Neurite Density Imaging (SANDI), which decomposes the measured dMRI signal into three main sources: extra-cellular space, neurite and soma where, if there is any sensitivity to the size of that compartment, the diffusion MRI signal in each of these compartments will have a time-dependence. This enabled both MR apparent soma size and soma signal fraction to be estimated non-invasively using dMRI for the first time.

However, the inverse problem of using complex multi-compartment models to infer microstructural information from the diffusion-weighted signal can be highly ill-posed and give rise to degeneracies in the model-parameter estimation, i.e., in the forward sense, completely different sets of model parameters predict the same dMRI signals (Jelescu et al., 2016; Novikov et al., 2018b; Jones et al., 2013; Lampinen et al., 2017, 2020, 2019). SANDI, like any other multi-compartment model (Jelescu et al., 2016), may suffer from the same degeneracy problems.

In SDE, the MR signal is sensitized to diffusion using a pair of gradient pulses that encode the position of the spins along the axis defined by the diffusion gradients. Double Diffusion Encoding (DDE) contains two pairs of pulsed-field gradients that are separated from each other with a mixing time *τ* (Cory et al., 1990; Callaghan, 2011). This approach has been utilized by several groups for extracting microstructural information (Özarslan et al., 2009; Jespersen et al., 2013; Benjamini et al., 2014; Ianuş et al., 2016; Yang et al., 2018b; Coelho et al., 2019). A framework called q-space trajectory imaging (QTI) was recently introduced by (Westin et al., 2016) to probe tissue using different gradient waveforms. The traditional, pulsed field gradient sequences attempt to probe a point in q-space but in q-space trajectory encoding, time-varying gradients are used to probe a trajectory in q-space, and the b-matrix is defined as an axisymmetric second order tensor (Topgaard, 2017). In this framework, SDE is just a special realization of linear tensor encoding (LTE) where the b-tensor has only one non-zero eigenvalue as all gradients are along the same axis. DDE is a special case of planar tensor encoding (PTE) as all gradients lie on a plane and the b-tensor has two non-zero eigenvalues. In spherical tensor encoding (STE) the gradients may point in all directions giving rise to a rank-3 b-matrix. Recently, b-tensor encoding has been used to resolve degeneracy problem (Reisert et al., 2019; Coelho et al., 2019; Fieremans et al., 2018; Gyori et al., 2019; Lampinen et al., 2020).

While these studies have shown improvement in the accuracy of parameter estimates, a consideration of time-dependence of the diffusion-weighted signal can be used as another feature to add information. In particular, Gyori et al. (Gyori et al., 2019) have recently proposed a method based on a three-compartment model to estimate neurite and soma features (e.g. signal fractions and intra-compartment apparent diffusivities) from combined LTE and STE data. Notably, Gyori et al. treated the signal coming from the spherical compartment as a simple monoexponential with a fixed, time-invariant small diffusivity. This implicitly assumes that the spherical compartment would show the same signal behavior for linear and spherical tensor encoding for a given b-value. However, Lundell et al. (Lundell et al., 2019) demonstrated that this is only true in restricted geometries when the LTE and STE waveforms have the same frequency power spectra.

In this study, we used b-tensor encoding with variable power spectra (including LTE and STE waveforms that were not spectrally matched to each other) to investigate whether and how it improves fitting of the SANDI model. Here, we exploit all three forms of b-tensor, i.e., LTE, STE, and PTE. We adopted van Gelderen’s model of the spherical compartment (Vangelderen et al., 1994), which explicitly includes both Δ and *δ*. The challenge in using the free gradient waveforms, however, is that Δ and *δ* are poorly defined, and so the time-dependency of the obtained signal is not well-defined in the time domain. Therefore, to find a closed-form for the diffusion-weighted signal decay in the spherical compartment, we adopted this model to the frequency domain (Nilsson et al., 2017; Stepišnik, 1993; Lundell et al., 2019).

The main findings of this paper are as follows:

- **Noise Sensitivity:** Even when complementing LTE with STE- and PTE-data, fitting the spherical pore properties remains challenging (when the signal fraction is small, i.e. < ~10%), with simulations showing biases in parameter estimates. Here we demonstrate that it is predominantly the Rician distribution of the noise (and associated noise floor) that impacts the estimation of spherical compartment properties. Such biases disappear when simulating purely Gaussian noise). However, if the Rician noise floor is accounted for in the model-fitting (albeit naively) – much of the noise-floor induced bias is ameliorated.
- **Lower Bound on Sphere Signal Fraction:** Here, by using the F-statistic to compare nested models (i.e., those that do or do not include a sphere fraction) in simulated data where the spherical pore properties are varied systematically, it was possible to identify a lower bound on the detectable MRI signal fraction limit. This was around 10% for data with SNR = 50.
- **Lower Bound on Spherical Pore Size:** The empirical lower limit on spherical pore size in brain tissue was estimated by comparing exponents in powerlaw relationships between the dMRI signal and b-value fitted to simulated data, with exponents observed empirically *in vivo*. We employed two different ways of establishing the lower bound for spherical radius estimates in white matter. The first, examining power-law relationships between the DW-signal and diffusion weighting in empirical data, yielded a lower bound of 7*μm*, while the second, pure Monte Carlo simulations, yielded a lower limit of 3*μm*. In addition, there is little differentiation in signal attenuation for low radii spherical pores (e.g. *R*_sphere_ < 4*μm*, Fig. .3 (a)).
- **Sticks + Ball + Sphere *vs* Cylinders + Ball + Sphere:** In addition to the challenge of estimating sphere size, the fitting becomes even more challenging if we have cylinders instead of sticks. As shown by Veraart et al. (Veraart et al., 2020), at 300 mT/m, and with appropriate diffusion times, we have sensitivity to the internal perpendicular diffusivity in cylindrical pores (demonstrated by a break from a power-law relationship between signal intensity and b-value). A challenge then arises when trying to disentangle two time-dependencies by varying the same experimental parameter (i.e, changing the frequency content of the gradient-encoding waveform). With currently-available pipelines, this prevents reliable estimates of spherical pore properties in white matter when there is sensitivity to intra-axonal radial diffusivity, and indeed may plague grey matter modelling if there is sensitivity to water in the astrocytic processes. We should note that most of the cellular projections are smaller than 3 microns in radius, while the majority of soma are above 3 microns (Savtchenko et al., 2018; Di Benedetto et al., 2016; Zhang et al., 2016; Papageorgiou et al., 2011; Fannon et al., 2015; Mohamed et al., 2020). Therefore, the ambiguity here is more relevant to WM voxels given a low soma density there.

## 2. Theory

Multi-compartment models express the diffusion-weighted signal as the sum of several compartments.

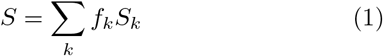

where *f_k_* is the signal fraction (∑_*k*_ *f_k_* = 1) and *S_k_* is the signal from the *k*th compartment.

For a general **B**-matrix, the diffusion-weighted MR signal is modeled as:

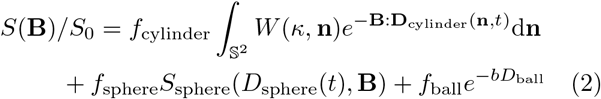

where *f*_cylinder_, *f*_ball_, *f*_sphere_, 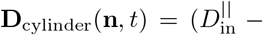 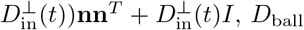, *D*_ball_ and *D*_sphere_ are the cylinder, ball and sphere signal fractions and diffusivities, respectively (Murday and Cotts, 1968). *W* (**n**) is the Watson orientation distribution function (ODF) and *κ* is the dispersion parameter. The diffusion weighting matrix **B** is given by 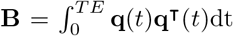 where 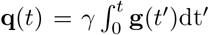 (Eriksson et al., 2015; Westin et al., 2016), and *γ* is the gyromagnetic ratio. Axial and radial elements in the diagonal axisymmetric b-tensor are *b*_||_ and *b*_⊥_ respectively, b-value, *b* is the trace of *B*-matrix and *b*_Δ_ = (*b*_||_ − *b*_⊥_)/*b*. For linear (LTE), planar (PTE) and spherical (STE) tensor encoding, *b*_Δ_ = 1, −1/2, and 0 respectively (Eriksson et al., 2015).

To remove the effect of fiber orientation dispersion (Jespersen et al., 2013; Lasič et al., 2014), the acquired signal is averaged over all diffusion directions for each shell. This so-called ‘powder-averaged’ signal (Callaghan et al., 1979; Edén, 2003) has less complexity than the orientation-dependent signal, and yields a signal whose orientationally-invariant aspects of diffusion are preserved but with an orientation distribution that mimics complete dispersion of anisotropic structures. Compartmental diffusion is represented with axisymmetric diffusion tensors which are described by isotropic diffusivity, *D*_I_ = 1/3*D*_||_ + 2/3*D*_⊥_, and anisotropy, *D*_Δ_ = (*D*_||_ − *D*_⊥_)/(*D*_||_ + 2*D*_⊥_) where *D*_||_ and *D*_⊥_ are the axial and radial diffusivities, respectively. *D*_Δ_ changes between −1/2 for a planar tensor to 1 for a stick. The signal attenuation from the *k*th compartment is given by (Lampinen et al., 2019; Eriksson et al., 2015):

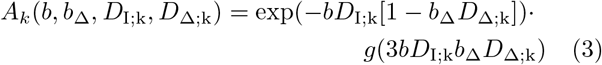

where

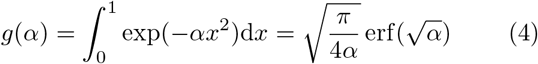

and erf(.) is the error function (Callaghan et al., 1979). Diffusion inside the sphere and ball is isotropic (*D*_Δ;sphere_ = 0, *D*_Δ;ball_ = 0) while for the cylinder and stick it is anisotropic (*D*_Δ;cylinder_ > 0, *D*_Δ;stick_ = 1). Therefore, the full signal equation is given by:

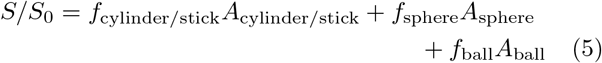

### 2.1. Two and three-compartment models

#### 2.1.1. Cylinder + Ball + Sphere (Extended SANDI Model)

In the original SANDI framework, the dMRI signal in brain tissue is assumed to arise from three main non-exchanging compartments: (i) intra-neurite (modeled as diffusion in sticks); (ii) intra-soma (modeled as diffusion constrained to a sphere); and (iii) extra-cellular (modeled as isotropic Gaussian diffusion). Here we extend this model to consider the perpendicular diffusivity in the intra-neurite compartment, 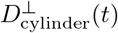, thereby modeling it with cylinders instead of sticks. We additionally explore the feasibility of modeling the intra-axonal perpendicular diffusivity and the additive spherical, *D*_sphere_(*t*), sensitivity simultaneously.

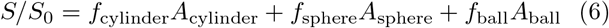

For complex gradient waveforms, the diffusion time is ill-defined. We therefore consider the diffusion spectrum 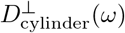, *D*_sphere_(*ω*) (Stepišnik, 1993; Lundell et al., 2019) in our analyses of compartment size.

The restricted DW-signal inside the sphere and cylinder is *S* = exp(−*ρ*) where *ρ* is (Stepišnik, 1993; Lundell et al., 2019):

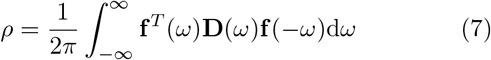

where 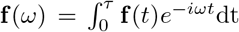, 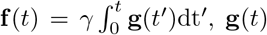 is the gradient waveform. **D**(*ω*) can be expressed with a rotation matrix **R** as **D**(*ω*) = **R**Λ(*ω*)**R**^−1^ where Λ(*ω*) is the diagonal matrix containing diffusion spectra *λ*_j_(*ω*) along the restriction principal axes. The analytical expression for *λ*_j_(*ω*) in the case of restricted diffusion in planar, cylindrical and spherical geometries, is the weighted sum of negative Lorentzians:

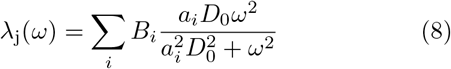

where for a cylinder, *λ*_1_(*ω*) = *λ*_2_(*ω*) = *λ*(*ω*) and *λ*_3_(*ω*) = 0 and

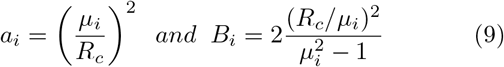

where *μ_i_* are the roots of 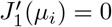 and 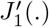 is the Bessel function of the first kind and order (Stepišnik, 1993; Nilsson et al., 2017; Lundell et al., 2019) and *R_c_* is the cylinder radius.

For a sphere, *λ*_1_(*ω*) = *λ*_2_(*ω*) = *λ*_3_(*ω*) = *λ*(*ω*) and

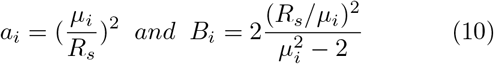

where *μ_i_* are the roots of the derivatives of the first order spherical Bessel function 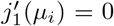 and *R_s_* is the sphere radius. *D*_0_ is fixed at 3*μm*^2^/*ms* for the sphere, as proposed in (Palombo et al., 2020) and 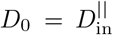 for cylindrical geometry.

#### 2.1.2. Stick + Ball + Sphere (R_cylinder_ = 0 (Original SANDI Model)

We define a three-compartment model, Stick + Ball + Sphere to investigate the sensitivity of the diffusion signal to the sphere radius and signal fraction. This model is the same as the original SANDI model with the difference that here we use b-tensor encoding and frequency-domain analysis.

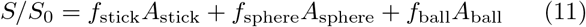

#### 2.1.3. Stick + Ball (Behrens et al., 2003) (f_sphere_ = 0 and R_cylinder_ = 0)

In this section, we compare the Stick + Ball + Sphere model with a Stick + Ball model and provide the range of sphere signal fractions and radii that make these two models significantly different. This latter model is the simplest model and does not have any time dependency.

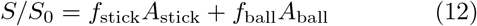

## 3. Method

Using both numerical simulations and *in vivo* experiments in healthy volunteers, we explore estimation of the soma size and density *R*_sphere_ and *f*_sphere_ using the combination of efficient gradient waveforms for LTE, PTE, and STE. Note: as we analyze the powder-averaged/orientationally-averaged signal, we do not need to estimate orientational dispersion. We study the challenges of the fitting landscape, the effect of noise, the lower limit on detectable signal fraction, the empirical lower limit on detectable pore size, and the challenge of disentangling two time-dependent properties (cylinder and sphere radius) of the model.

### 3.1. Fitting landscape

There is little differentiation in signal attention for low radii (*R_s_* < 4*μm*) (Fig. .3(a)). When there is low sensitivity to some parameters, the numerical optimization algorithm terminates prematurely and therefore the estimates are not accurate.

### 3.2. Noise sensitivity

To explore the sensitivity of parameter estimation to noise perturbations, we simulated three different scenarios: (i) addition of Gaussian noise to the magnitude of the signal; (ii) addition of Gaussian noise to the real and imaginary channels which results in Rician-distributed magnitude signal; and (iii) addressing the noise-floor problem in case (ii) with a (simple) correction. In general, when there is Gaussian noise in the signal, averaging improves the signal to noise ratio (SNR) and because of the orientational-averaging used in this work, we expect some improvement in the SNR in the first scenario and therefore better estimates of the model parameters. In the second scenario, the signal is corrupted by Rician-distributed noise, and therefore the orientational-averaging that improved the SNR in case (i), does not remove the non-zero positive ‘noise-floor’ bias in Rician-distributed noise and therefore we expect some bias in the parameter estimates. In the third scenario, we use a simple correction for the Rician bias in case (ii). To estimate the standard deviation of the noise, we include the noise floor in the model so that the predicted signal is 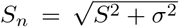 where *S* is our original model prediction and *S_n_* is the prediction after accounting for the noise floor (Jones and Basser, 2004; Koay et al., 2009; Pieciak et al., 2016b, 2018, 2016a). We expect some improvement in the parameter estimates in case (iii) compared to case (ii) but the results of case (i) are expected to best out of all three cases.

### 3.3. Lower Bound on Resolvable Spherical Pore Signal Fraction

Here we considered that the framework had sensitivity to the sphere signal fraction if the inclusion of a sphere component to a stick + ball model was statistically supported by an F-test. To determine the lower bound on the spherical signal fraction and the pore size that can be detected using a diffusion-weighted signal, we systematically varied both parameters, while comparing the fit from two models: (i) the stick + sphere + ball model (three-compartments <u>including</u> a spherical component); and (ii) a stick + ball model (two-compartment <u>without</u> a spherical component). To test whether inclusion of the spherical compartment was needed to describe the signal (thereby showing sensitivity to this component) we considered the stick + sphere + ball model justified if the p-value from the F-test was less than 0.05 (Nilsson and Alexander, 2012; Panagiotaki et al., 2012). Here, the F-statistic is calculated as F = (SSR_1_ − SSR_2_)(N − M_2_)/(SSR_2_(M_2_ − M_1_)) where SSR is the sum of squared residuals, M is the number of fitted parameters of the simplified (1) and full SANDI model (2), and N is the number of measurements. The p-value is estimated using p = 1 fcdf(F, M_2_ − M_1_, N − M_2_) where fcdf is the cumulative distribution function of F-distribution.

### 3.4. Stick + Ball + Sphere vs Cylinder + Ball + Sphere

If ultra-strong gradients and diffusion-time settings are such that we do, indeed, have sensitivity to the intra-neurite perpendicular diffusivity, then the intra-neurite compartment should be more correctly modeled using cylinders instead of sticks (Veraart et al., 2020). This introduces an additional challenge, as we now have two compartments (sphere and cylinder) with a diffusion-time dependence. To explore this, we conducted further simulations to investigate the impact of including a non-zero perpendicular intra-neurite diffusivity (or, rather, a cylinder with a finite radius of *R*_cylinder_ = 4 *μm*) on estimation of spherical pore-size.

### 3.5. Empirical Lower Bound on Spherical Pore Size

To identify the empirical lower bound on spherical pore size, we simulated signals for experiments with fixed diffusion time, Δ = 37.05*ms*, and *δ* = 29.65*ms* (matching our *in vivo* experimental set-up), diffusivities 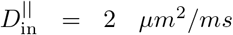 and *D*_ball_ = 2 *μm*^2^/*ms*, but with variable sphere signal fractions *f*_sphere_ = (0.01 : 0.01 : 0.1, 0.15, 0.2 : 0.1 : 1), *f*_ball_ = *f*_cylinder_ = (1 − *f*_sphere_)/2, and sizes, *R*_sphere_ = 1 : 0.5 : 10 *μm*. For each set of signals, we fitted a power-law (Veraart et al., 2019; McKinnon et al., 2017; Lampinen et al., 2017) to the direction-averaged signal from the LTE measurements for *b* = 6, 7.5, 9, 10.5*ms/μm*^2^ according to (*S/S*_0_ = *βb*^−*α*^) and then compared the values of the exponent, *α*, with values observed empirically *in vivo* to establish a lower bound on the size of spherical pores. The rationale behind the choice of using the power-law to drive an empirical conclusion is that it is free of any model assumptions, and simply considers the rate of signal decay versus b-value. We know that the *α* value for a pure stick-like geometry is 0.5, and thus any deviation from this value is indicative of sensitivity to an additional compartment (a deviation from the stick-like geometry could be due to any shape that is not stick-like). The compartment that we choose to change is, in fact, a spherical compartment. By systematically increasing the size of the spherical compartment until such a deviation is detected, we can obtain an empirical lower bound on the spherical compartment. Any sensitivity to the intra-axonal perpendicular diffusivity would make the signal decay faster (Veraart et al., 2020) but with the timing parameters used here, we do not expect any such sensitivity.

### 3.6. Simulations

The numerical simulations were performed using the model in Eq (2), with *f*_sphere_ = 0.01 : 0.01 : 0.1, 0.15, 0.2 : 0.1 : 1, *f*_ball_ = *f*_stick_ = (1 − *f*_sphere_)/2, 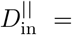 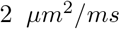, *D*_ball_ = 0.6 *μm*^2^/*ms*, *R*_sphere_ = 1 : 0.5 : 10 *μm*, and *R*_cylinder_ = 4 *μm*. The simulated protocol matched the *in vivo* protocol and comprised 10 *b* = 0 and 8 non-zero shells (*b* = 1, 2, 3, 4.5, 6, 7.5, 9, 10.5 *ms/μm*^2^) in (10, 31, 31, 31, 31, 61, 61, 61, 61) directions for LTE and 5 shells (*b* = 1, 2, 3, 4.5, 6 *ms/μm*^2^) in (31, 31, 31, 31, 61) directions for PTE and 5 shells for STE (*b* = 0.2, 1, 2, 3, 4.5 *ms/μm*^2^) in (6, 9, 9, 12, 15) directions and SNR = 50 with Rician noise. The 61 and 31 directions were optimized based on (Knutsson, 2018). The noisy diffusion signal was modeled according to the following:

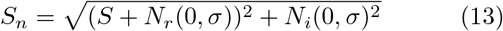

where *S_n_* and *S* are the noisy and noise-free signals, respectively, and *N_r_* and *N_i_* are the normal distributed noise in the real and imaginary images respectively with a standard deviation of *σ* (Aja-Fernández and Vegas-Sánchez-Ferrero, 2016; Jones and Basser, 2004). Here SNR level is defined as 1/*σ*. For each b-tensor shape, and for each b-value, the diffusion signal was averaged over all directions in a shell.

### 3.7. In vivo *Data*

Two healthy participants who showed no evidence of a clinical neurologic condition were scanned with the approval of the Cardiff University School of Psychology Ethics Committee. Magnetization-prepared rapid gradient echo (MPRAGE) images were also acquired for anatomical reference. 192 sagittal slices with TE = 2.3 ms, TR = 1900 ms, TI = 900 ms, and a voxel size of 1 mm isotropic and 256 × 256 matrix size were acquired in 5 minutes.

Diffusion-weighted images were acquired with the protocol detailed in the simulation section 3.6 on a 3T Connectom MR imaging system with 300 mT/m gradients (Siemens Healthineers, Erlangen, Germany). Forty-two axial slices with 3*mm* isotropic voxel size and a 78 × 78 matrix size, TE = 88 ms, TR = 3000 ms, partial Fourier factor = 6/8, and heat dissipation limit = 1, were obtained for each individual. The total acquisition time was around one hour. To take full advantage of q-space trajectory imaging, it is imperative to respect the constraints imposed by the hardware, while at the same time maximizing the diffusion encoding strength. Sjolund et al. (Sjölund et al., 2015) provided a tool for achieving this by solving a constrained optimization problem that accommodates constraints on maximum gradient amplitude, slew rate, coil heating, and positioning of radiofrequency pulses. The gradient waveform is optimized and Maxwell-compensated (Szczepankiewicz et al., 2019) based on a framework that maximizes the b-value for a given measurement b-tensor and echo time. Substantial gains in terms of reduced echo times and increased signal-to-noise ratio can be achieved, in particular as compared with naive planar and spherical tensor encoding. Duration of the first, pause, and the second waveform in Fig. .2 were [29.6, 7.4, 29.6] *ms* for LTE and [35.6, 7.4, 28.6] *ms* for PTE and STE. The slew rate was 13.8, 62, and 51.1 mT/m/ms for LTE, PTE, and STE, respectively.

### 3.8. Preprocessing

The diffusion weighted images were corrected for Gibbs ringing (Kellner et al., 2016). We acquire some interleaved b0 images between the diffusion-weighted images (DWIs) to use for motion correction. In PTE and STE data, we registered the interleaved b0 images to the first b0 image and used the corresponding transformation to correct the motion in the DWIs. In LTE data the eddy current and subject motion were corrected by FSL EDDY (Andersson and Sotiropoulos, 2016) and finally the gradient nonlinearity was corrected by the method proposed by Rudrapatna et al. (Rudrapatna et al., 2020, 2018). We applied a 3D Gaussian filter with a standard deviation of 0.5 to the preprocessed data to make the images smooth. We normalized the direction-averaged signal based on the b = 0 *s/mm*^2^ signal in each voxel.

### 3.9. Regions of interest

We defined five regions of interest (ROIs) in the splenium and internal capsule as white matter regions and putamen, ventrolateral thalamus, and mediodorsal thalamus, as gray matter regions. To define anatomical regions of interest, publically-available atlases were co-registered to each participant’s T_1_-weighted MPRAGE image. The white matter ROIs were obtained from the JHU atlas (Mori et al., 2005), while the putamen ROI was obtained from the work of Tziorti et al. (Tziortzi et al., 2011) and ventrolateral thalamus and mediodorsal thalamus from (Danos et al., 2003). The MPRAGE image was co-registered to the diffusion-weighted data, and the resulting transform applied to the ROIs to translate them to the native diffusion-weighted space. The five ROIs are illustrated in Fig. .1.

**Figure .1:**
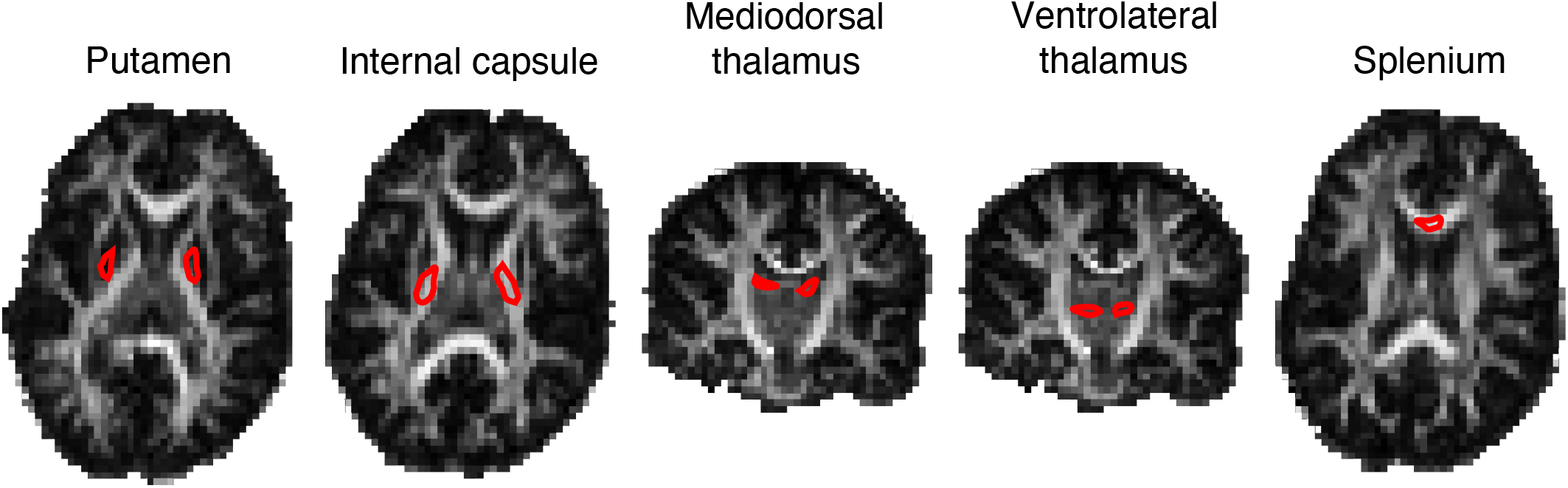
Location of the five ROIs used for the quantitative analysis of this study overlaid on the FA image of one subject. The posterior limb of the internal capsule, splenium, putamen, ventrolateral thalamus, and mediodorsal thalamus are illustrated as red ROIs on the FA map.

**Figure .2:**
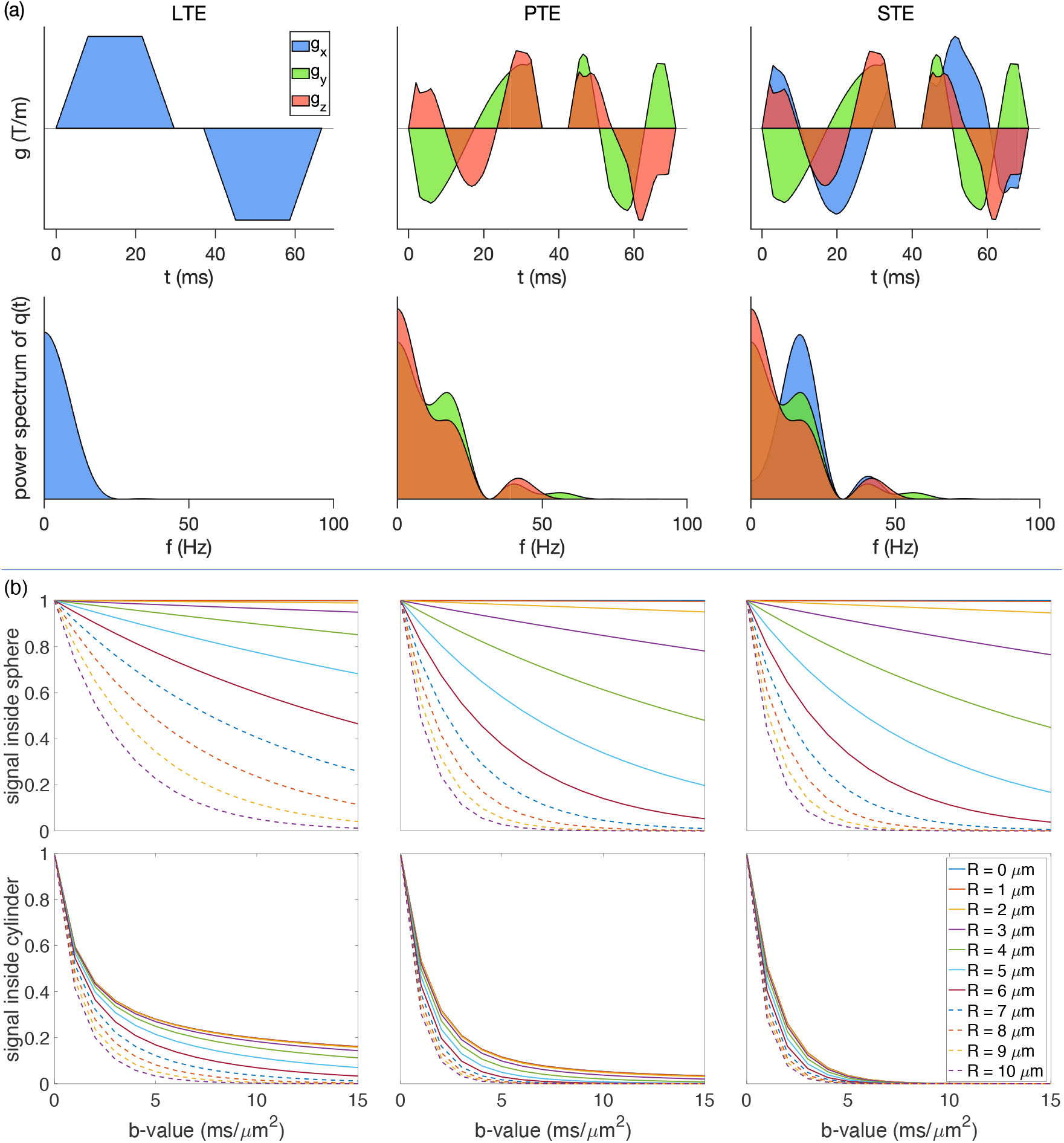
(a) The free gradient waveforms of the linear, planar, spherical tensor encoding and the corresponding frequency power spectra. (b) The signal decay inside the spherical and cylindrical compartments using different encoding schemes.

**Figure .3:**
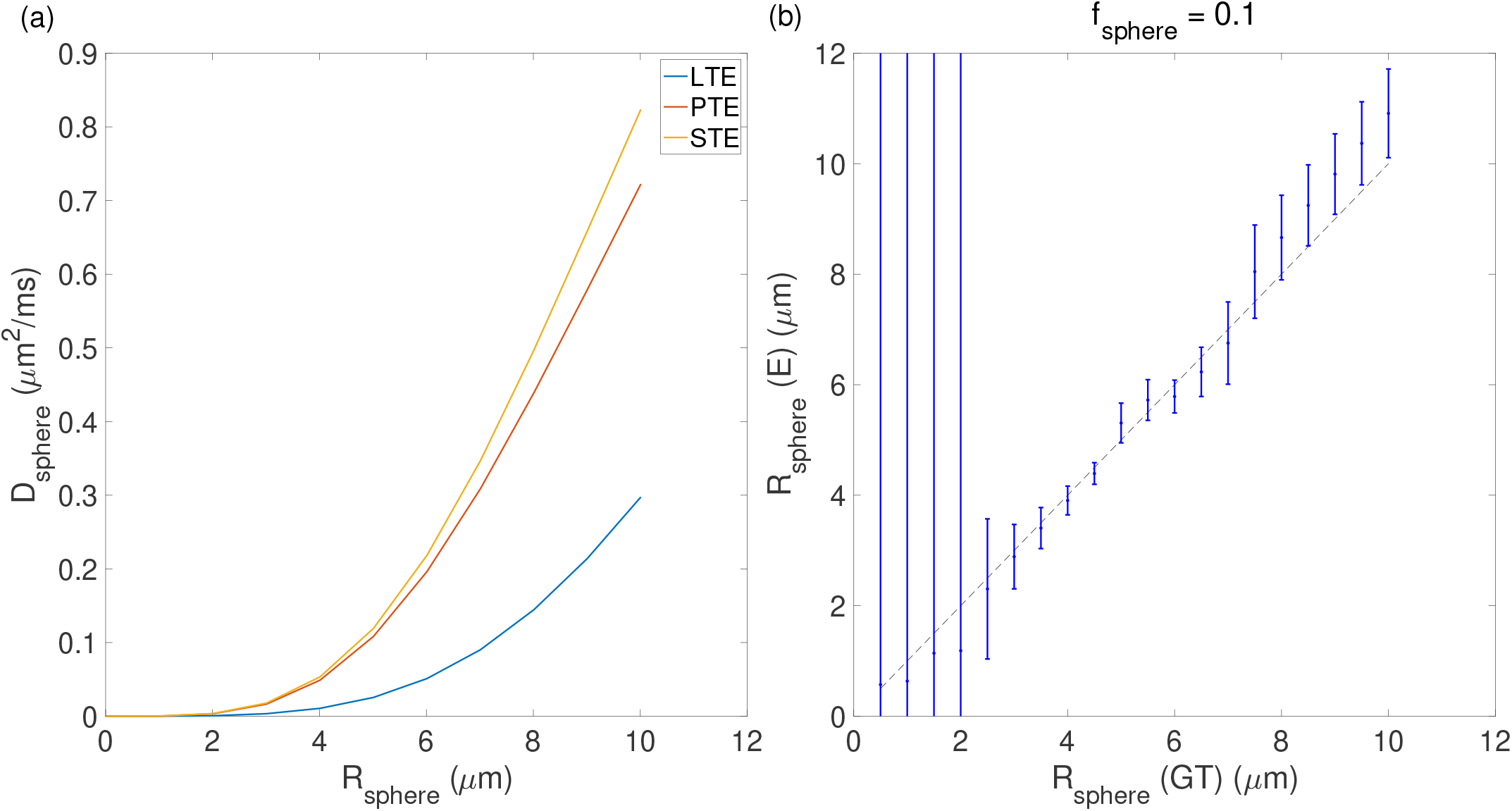
(a) The changes in the apparent diffusivity (*D*_sphere_) *versus* the radius of the sphere (*R*_sphere_) for linear, planar and spherical tensor encoding (LTE, PTE, and STE), (b) the results of sphere size estimation (The error bars show the confidence interval, dashed black line is the line of identity, GT = ground truth and E = estimated).

### 3.10. *Goodness of fit for* in vivo *data*

To check the stability of the model fit and that the global minima of the cost function had been found, we first fixed the signal fraction of the spherical compartment and estimated the remaining parameters (The same procedure was used by Lampinen et al. (Lampinen et al., 2019) for finding the stick fraction that can be detected reliably). To assess the precision of parameter estimation, a metric of the goodness-of-fit (see below) was plotted for different values of sphere signal fraction, which was varied systematically between zero and one in 40 equal steps. If the model determined all parameters unequivocally, a clear optimum in the goodness-of-fit would be seen for some signal fractions. Conversely, a flat plot of the goodness-of-fit over a wide range of signal fractions would indicate degeneracy in the fitting, i.e. two or more sets of solutions yield a similarly good fit. For each ROI, all the voxels contained therein were concatenated to provide a sufficient number of data points for our estimation.

Goodness-of-fit was determined using reduced chi-square, 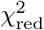 (Andrae et al., 2010). The reduced chi-square or normalized residual variance (NRV) (Lampinen et al., 2019) was obtained by dividing the residual variance 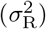 by the variance of noise 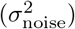:

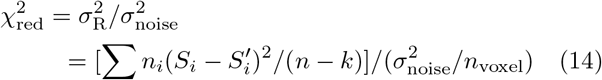

where *n_i_* is the number of directions for *i*th *b* and *b*_Δ_. *S_i_* and 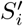 are the direction-averaged measured and predicted signals, *n* is the number of samples and *k* is the number of free parameters in the model. The standard deviation of the noise 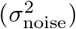 was estimated for each ROI.

## 4. Results

### 4.1. Simulations

Fig. .2 (a) shows the gradient waveforms used for the linear, planar and spherical tensor encoding and their corresponding frequency spectra. Fig. .2 (b) shows the signal decay inside the spherical and cylindrical compartments using different encoding schemes. It is important to note that, for a given b-value, STE results in the most signal loss, followed by PTE and then LTE. LTE appears relatively insensitive for small radii spherical compartments.

#### 4.1.1. Fitting landscape

Fig. .3 (a) shows the changes in apparent diffusivity of the spherical compartment (*D*_sphere_) as a function of radius of the spherical compartment for the three b-tensor shapes. Clearly, there is a large difference in the sensitivity to pore size, with LTE being the least sensitive and PTE and STE tracking each other closely in the plot of *D*_sphere_ *vs R*_sphere_. Notably, for all wave-forms, there is little differentiation in sphere signal attenuation for low radii, (e.g. *R*_sphere_ < 4*μm*). For small radii, the noise dominates over measurable effects. The error bars in Fig. .3 (b) show the confidence interval. In the fitting, we use *lsqcurvefit* in MATLAB which returns the jacobian at the solution, and then *nlparci* command is used to find the 95% confidence interval.

#### 4.1.2. Noise sensitivity

Fig. .4 shows the results of fitting the sphere radius for different sphere signal fractions under different noise simulations. The figure also shows the p-value of the F-test between the two and three-compartment models in the presence of Gaussian, Rician, and corrected Rician noise. Here, we take *p* < 0.05 as an indication that the full model (three-compartment) is preferred over the simplified model (two-compartment). When the sphere radius or the signal fraction of the sphere is small (*R*_sphere_ < 2*μm* and *f*_sphere_ < 0.05) the simplified model is preferred. Fig. .4 shows that the estimates are largely positively biased in the Rician-noise data, whereas their errors are symmetrically-distributed about the line of identity in Gaussian only. After Rician noise correction, it looks more like the Gaussian - in terms of symmetrical distribution around the line of identity.

**Figure .4:**
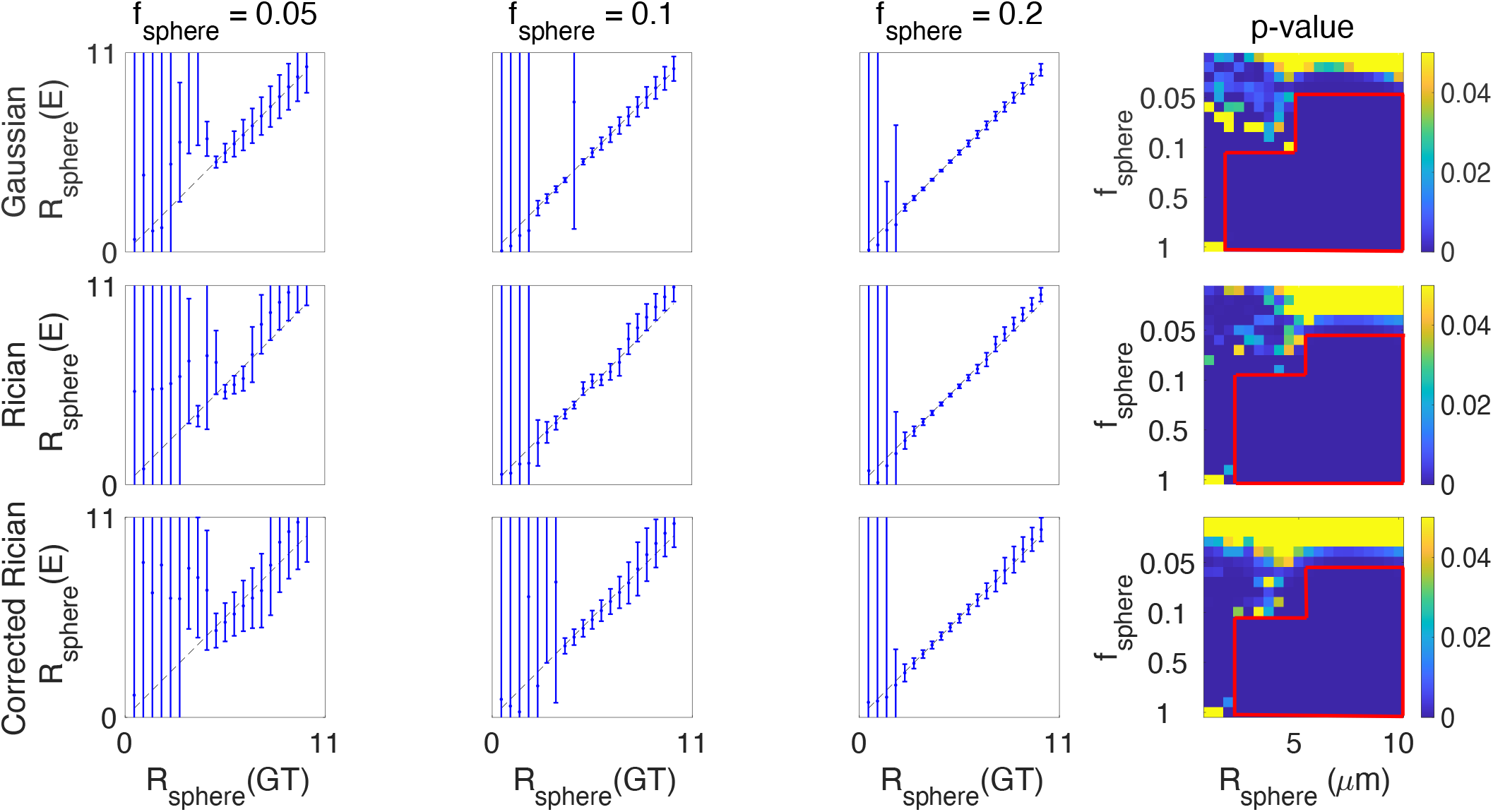
The results of fitting the sphere radius for different sphere signal fractions (GT = Ground Truth and E = Estimated). The figure also shows the p-value of the F-test in the presence of Gaussian, Rician, and corrected Rician noise respectively. The red rectangles in the right side plots show the areas that the three-compartment model is significantly different from the two-compartment model. The diagonal black line is the line of identity.

#### 4.1.3. Signal fraction resolution limit

The examination of the F-test measures indicates a lower bound on the signal fraction that can be detected or reasonably modeled from the diffusion-weighted signal. This is around 10% for SNR = 50 but this limit changes at different noise levels (Fig. .4). The changes of sphere radius estimates versus ground truth for a wider range of SNR values are shown in this link: https://bit.ly/StickBallSph. The lower limit scales inversely with SNR, i.e. as SNR increases, smaller sizes are detectable.

#### 4.1.4. *Stick + Ball + Sphere* vs *Cylinder + Ball + Sphere*

In addition to the challenge of estimating sphere size (given the above 4 points), the fitting becomes more challenging if we have cylinders instead of sticks because in this model the time-dependence of the signal can now arise from two independent sources (the cylinder and the sphere). Fig. .5 shows the estimated sphere size when there is a non-zero diameter cylinder, (SNR = 200). For signal fractions smaller than 0.2 and sphere radius larger than 5 microns, the estimated sizes deviate from the ground truth. This behavior is observed where the noise floor is very low (SNR = 200) in the simulated signal. The deviation from the ground truth in the presence of noise, for different noise levels, is shown in the following link: https://bit.ly/CylBallSph.

**Figure .5:**
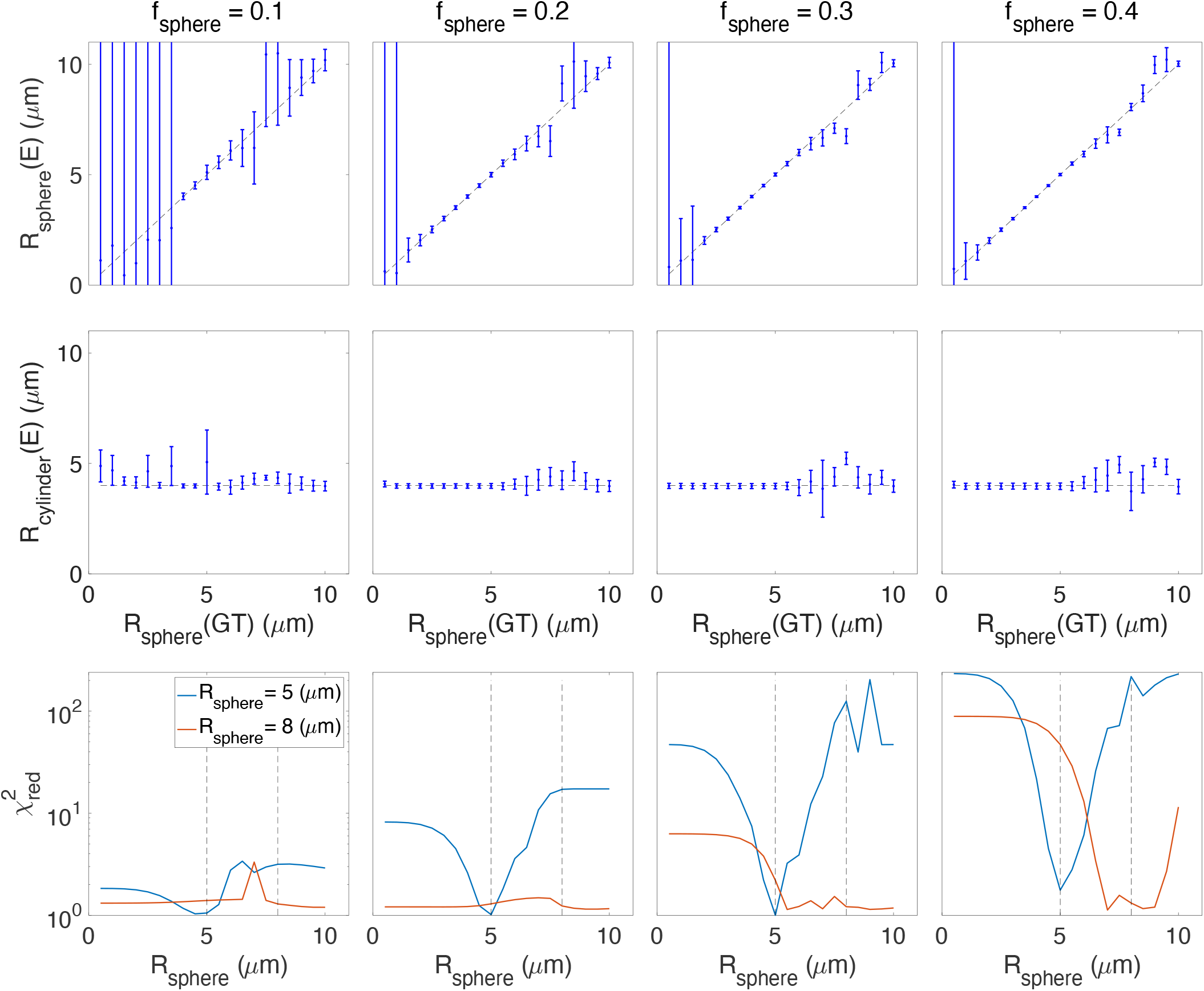
Estimated sphere and cylinder radii versus the ground truth sphere radius values for cylinder + ball + sphere model without noise (*SNR* = 200). The third row shows the reduced chi-square, 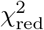, values for two scenarios where the sphere radius is fixed to 5 and 8 *μm*, blue and red curves respectively. Note that there is a sharp minimum for *R* = 5*μm* while the 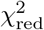 curve has several local minima or even flat in the *R* = 8*μm* case (GT = ground truth and E = estimated).

#### 4.1.5. Empirical lower limit on pore size

In addition to the lower bound imposed by the noise floor on the sphere signal fraction and size, there is an empirical lower limit on the pore size which can be derived from power-law measurements. Fig. .6 shows the effect of sphere signal fraction and size on the estimated exponent, *α*, in the power-law fit (*S/S*_0_ = *βb*^−*α*^) experiments. As will be discussed below, in our *in vivo* data, we observe power-law exponents greater than 0.5 in the white matter. Fig. .6 shows that to observe such values of *α* the sphere radius needs to be larger than 7 microns. We will discuss the implications of this key result in the context of the *in vivo* data below.

**Figure .6:**
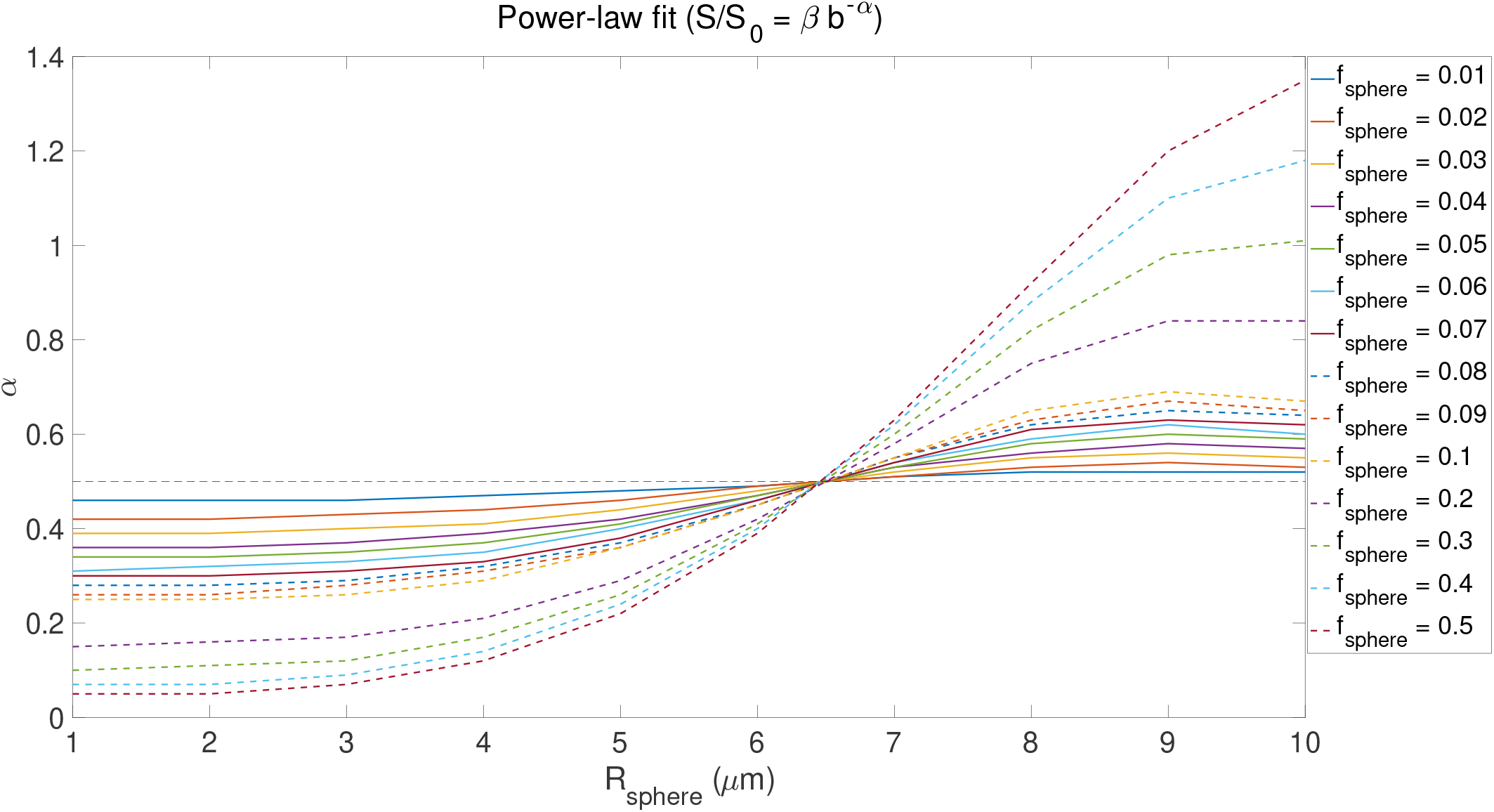
The effect of sphere size and signal fraction on exponent *α* (similar to Fig. 2 in (Palombo et al., 2018a)). (*f*_sphere_ = 0.01 : 0.01 : 0.1, 0.2 : 0.1 : 0.5, *f*_ball_ = *f*_stick_ = (1 − *f*_sphere_)/2, 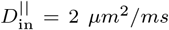, *D*_ball_ = 2 *μm*^2^/*ms*, *R*_sphere_ = 1 : 1 : 10 *μm*, *δ* = 29.65 *ms*, and Δ = 37.05 *ms*).

A good ‘sanity check’ is that after reconstructing the signal from the estimated parameters the exponent *α* in the power-law fit (*S/S*_0_ = *βb*^−*α*^) of the original and the modeled signal should not change considerably (see the last row of Fig. .7), otherwise, the parameters are representing the signal incorrectly. This approach has several limitations that should be acknowledged; The first limitation is that we used a pre-determined and fixed set of parameters in the simulation. If we had used a smaller diffusivity for the extra-cellular compartment, for the range of b-values used here (6 < *b* < 10.5 *ms/μm*^2^), there might have been some residual contribution from the extracellular compartment and therefore the exponent *α* and the behaviour of the signal would change. The second limitation is that we fixed the intra-neurite and extracellular signal fractions to be equal which may not reflect reality. Despite these limitations, we consider the result a useful empirical benchmark because the fixed diffusivities used in this experiment are close to the range of diffusivities estimated in previous works (Dhital et al., 2019). As such, the contribution from the extracellular compartment at high b-values will be negligible meaning that the only remaining signal contributions come from spheres and sticks.

**Figure .7:**
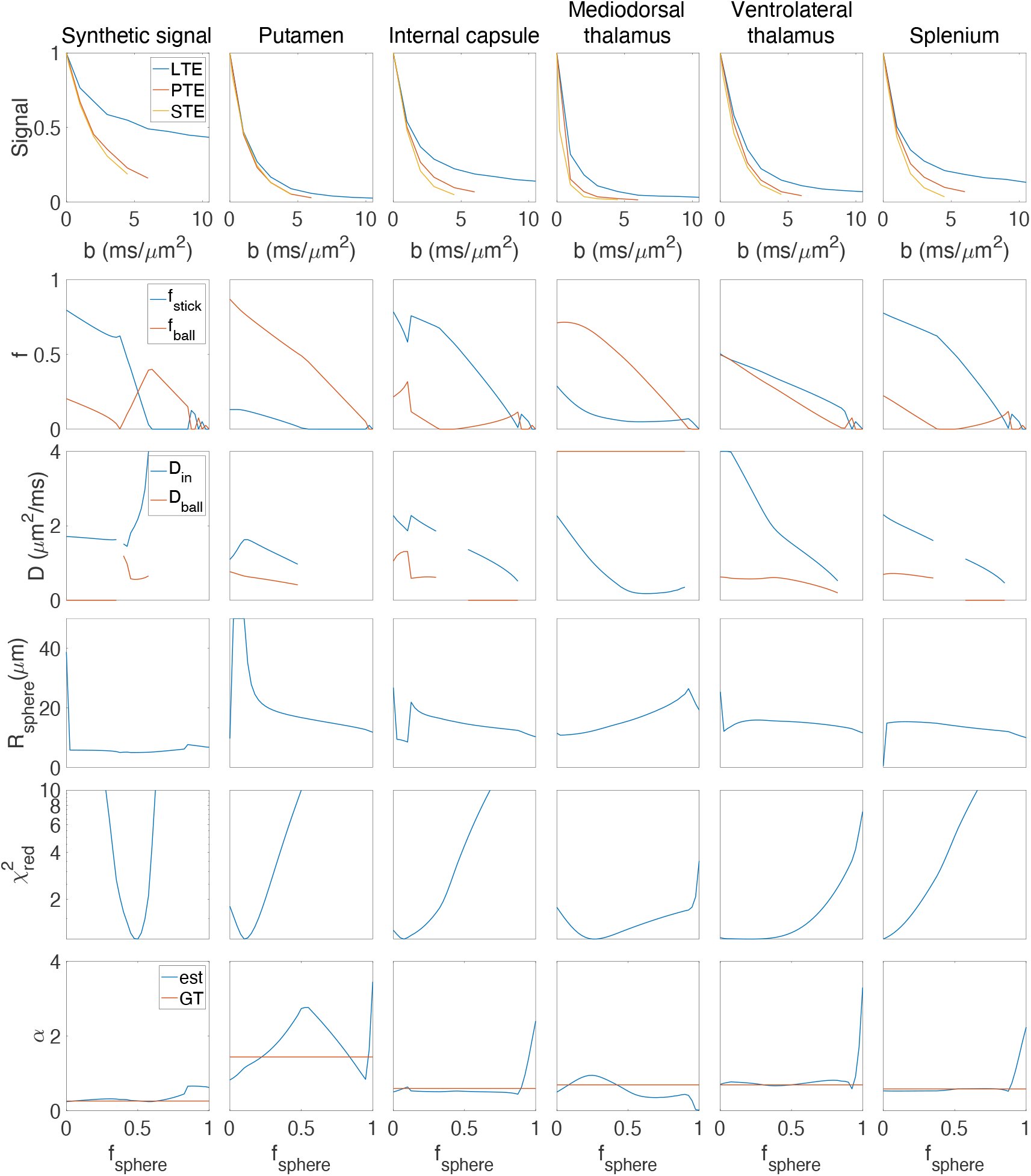
The results of fitting the stick + ball + sphere model to the diffusion-weighted signal by fixing the sphere signal fraction to different values. Five different ROIs of the brain are used here; putamen, internal capsule, mediodorsal thalamus, ventrolateral thalamus, and splenium. The mean value of the direction-averaged signal for each ROI is represented in the first row (in different columns). The second row shows the estimated signal fraction of stick (*f*_stick_) and ball (*f*_ball_) for different predefined sphere signal fractions (We used *f* as y-label here to show the signal fraction of both ball and stick in one plot.). The third row illustrates the parallel diffusivity of the stick 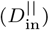 and the diffusivity of the ball (*D*_ball_) for different ROIs. The estimated radius of the sphere is illustrated in the fourth row. And finally, the last two rows show how well this model can explain the signal for different predefined sphere signal fractions (*f*_sphere_) in terms of reduced chi-square and power-law. The first column in the figure shows the results of fitting a synthetic signal. Note that we do not estimate diffusivity of the compartment when the signal fraction is estimated as zero, this is the reason for discontinuity in the plots of estimated diffusivities.

### 4.2. In vivo *results*

Here we provide *in vivo* results from fitting the Stick + Ball + Sphere model. Fig. .7 shows the results of fitting the model to the signal by fixing the sphere signal fraction to different values. Results are shown for five different ROIs: Splenium; internal capsule; mediodorsal thalamus; ventrolateral thalamus; and putamen. Note that, for the fitting, the data points from all three voxels in the ROI are concatenated to provide enough data points for a stable fit, but in the figure, the average of these three voxels is shown. The flat valleys in 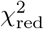 correspond to plausible sphere signal fractions in different ROIs of the brain, for example, 0-0.125 for Splenium and Internal Capsule and 0.2-0.3 for the Putamen (Fig. .7). When the valley is flat for a large range of sphere signal fractions, it means the data do not provide a unique solution and a range of parameters can represent the signal equally well. This may be related to a large range of acceptable parameters as shown by the second to the fourth row of Fig. .7. For example, acceptable 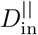 values for the stick compartment ranged between 2.2 and 2.5 *μm*^2^/*ms* in the splenium and between 1.2 and 1.5 *μm*^2^/*ms* in the putamen. If there was no degeneracy in the estimation of parameters and the data could provide useful information about the underlying microstructure, then the plot would have a sharp valley at the local optimum. Among the five ROIs we selected in this experiment, we see a quite sharp minimum for the putamen and mediodorsal thalamus (with *f*_sphere_ around 0.2-0.3), and the minima for white matter (splenium and internal capsule) clearly puts the sphere fraction in the sub 10% regime, which is where it is expected. The first column of Fig. .7 shows the parameter estimates for a synthetic signal generated with the following parameters; *f*_sphere_ = 0.5, *f*_ball_ = *f*_stick_ = 0.25, 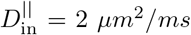, *D*_ball_ = 0.6 *μm*^2^/*ms*, *R*_sphere_ = 5 *μm*, and SNR = 100, Rician distributed signal. There is a sharp minimum in 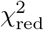 which shows there is only one set of parameters that fits the signal accurately.

Fig. .8 shows estimated parameter maps *in vivo*. When fitting on a voxel by voxel-level, the fitting is unstable and the resulting maps are not smooth. Besides, the large voxel size (3 mm) used here and the resulting problems with partial volume may affect the estimated parameters as well. Maps of the estimated parameters in Fig. .8 show a reasonable contrast that matches the results in (Palombo et al., 2020). The *f*_stick_ map has higher values in white matter tracts in the brain while *f*_sphere_ values are higher in the gray matter. The *f*_stick_ values in cortical GM range from 0.1 to 0.2 which is in agreement with recent works on estimating neurite density in GM using b-tensor encoding (Lampinen et al., 2017, 2019).

**Figure .8:**
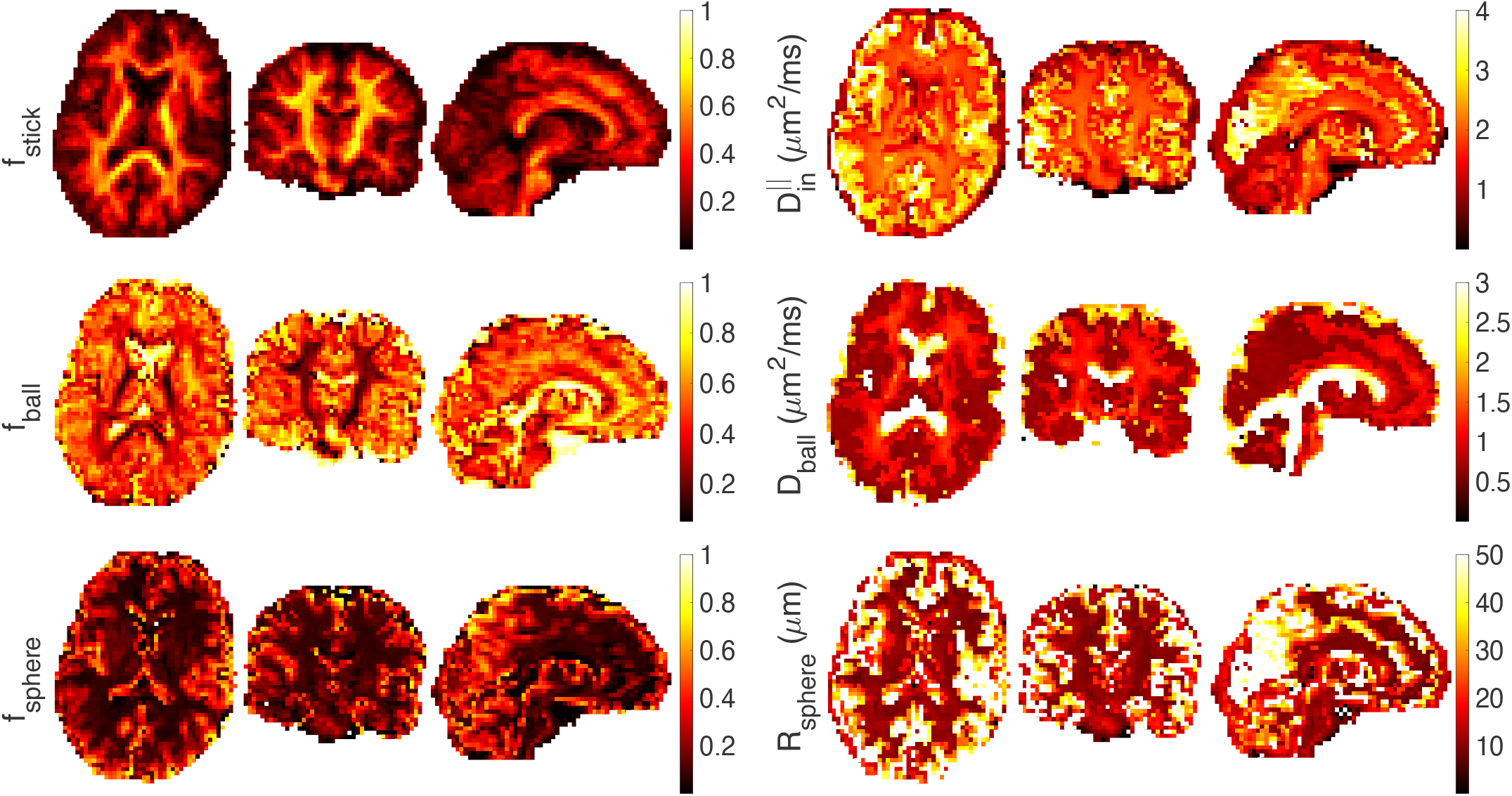
Estimated stick (*f*_stick_), ball (*f*_ball_), and sphere (*f*_sphere_) signal fractions, intra-axonal parallel diffusivity 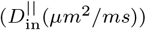, extra-cellular diffusivity (*D*_ball_(*μm*^2^/*ms*)), and sphere radius (*R*_sphere_(*μm*)) on axial, sagittal, and coronal views of the smoothed brain image (A 3D Gaussian kernel with standard deviation of 0.5 is used for smoothing).

Fig. .9 illustrates the estimated parameters of the ball+stick+sphere model including the noise floor as an extra parameter to fit.

**Figure .9:**
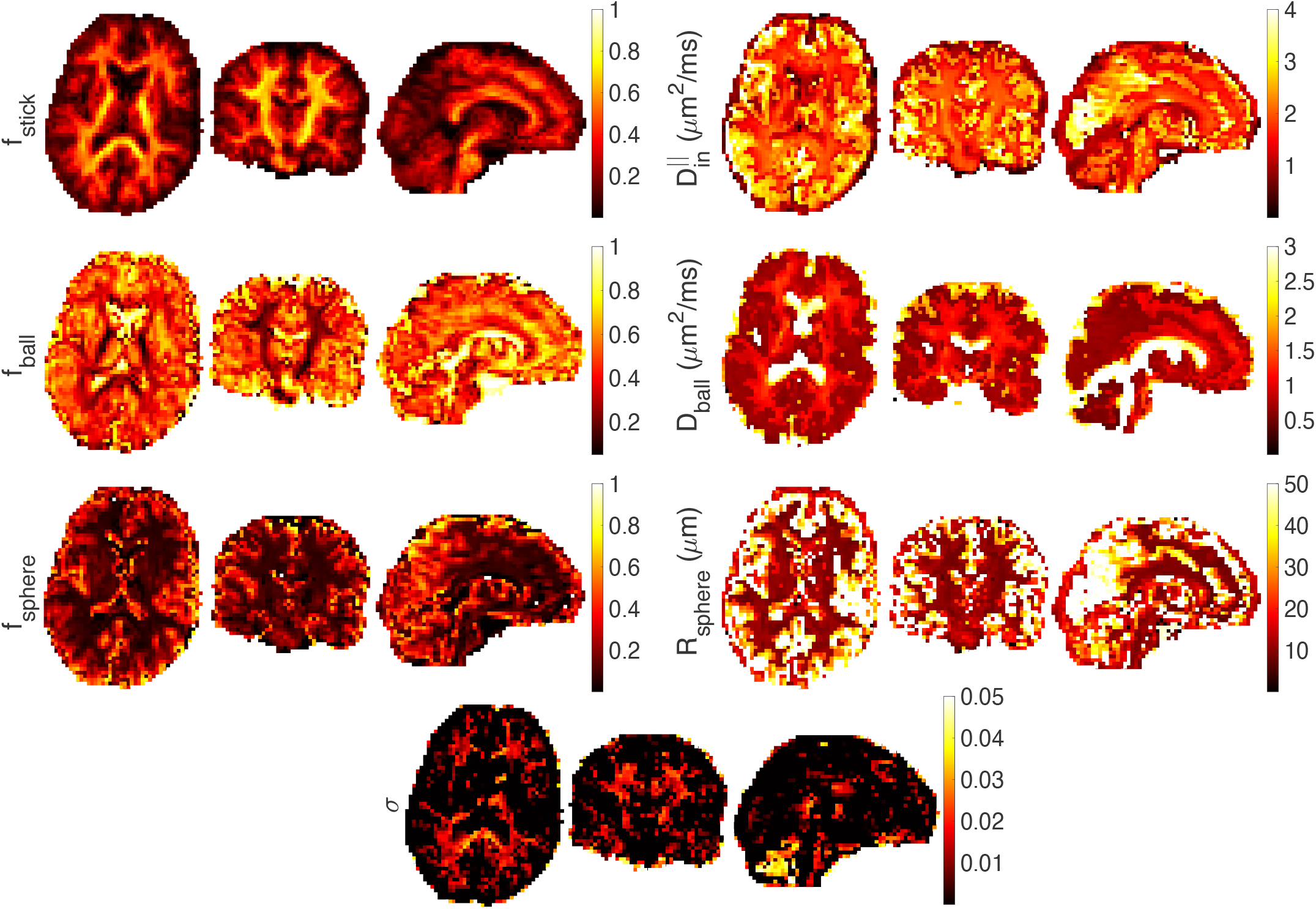
Estimated stick (*f*_stick_), ball (*f*_ball_), and sphere (*f*_sphere_) signal fractions, intra-axonal parallel diffusivity 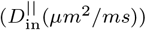, extra-cellular diffusivity (*D*_ball_(*μm*^2^/*ms*)), sphere radius (*R*_sphere_(*μm*)), and standard deviation of the noise (*σ*) on axial, sagittal, and coronal views of the smoothed brain image (A 3D Gaussian kernel with standard deviation of 0.5 is used for smoothing).

Fig. .10 shows the results of fitting a power-law (*S/S*_0_ = *βb*^−*α*^) to the diffusion-weighted signal. CSF has the fastest decay and therefore no signal remains from CSF in the high b-value data and the alpha is close to zero. The decay in the GM is faster than the WM and therefore the estimated *α* values are correspondingly larger in GM than WM. Within WM, *α* is usually larger than 0.5. As noted by Veraart et al. (Veraart et al., 2019) and McKinnon et al. (McKinnon et al., 2017), if there is only a stick compartment, then *α* = 0.5. The decay faster than the stick model may come from the exchange between compartments (Stanisz et al., 1997), sensitivity to the axon diameter (Veraart et al., 2020) or the presence of a non-negligibly-sized third compartment (Palombo et al., 2020) that makes the signal decay faster than 0.5. If the radius of the spherical compartment is less than approximately 7 microns then the exponent is smaller than 0.5. As will be seen in the map of *α* in Fig. .6, we do not observe values of *α* less than 0.5 in the white matter. This places a lower bound on the size of spherical pores in white matter of around 7 *μm*.

**Figure .10:**
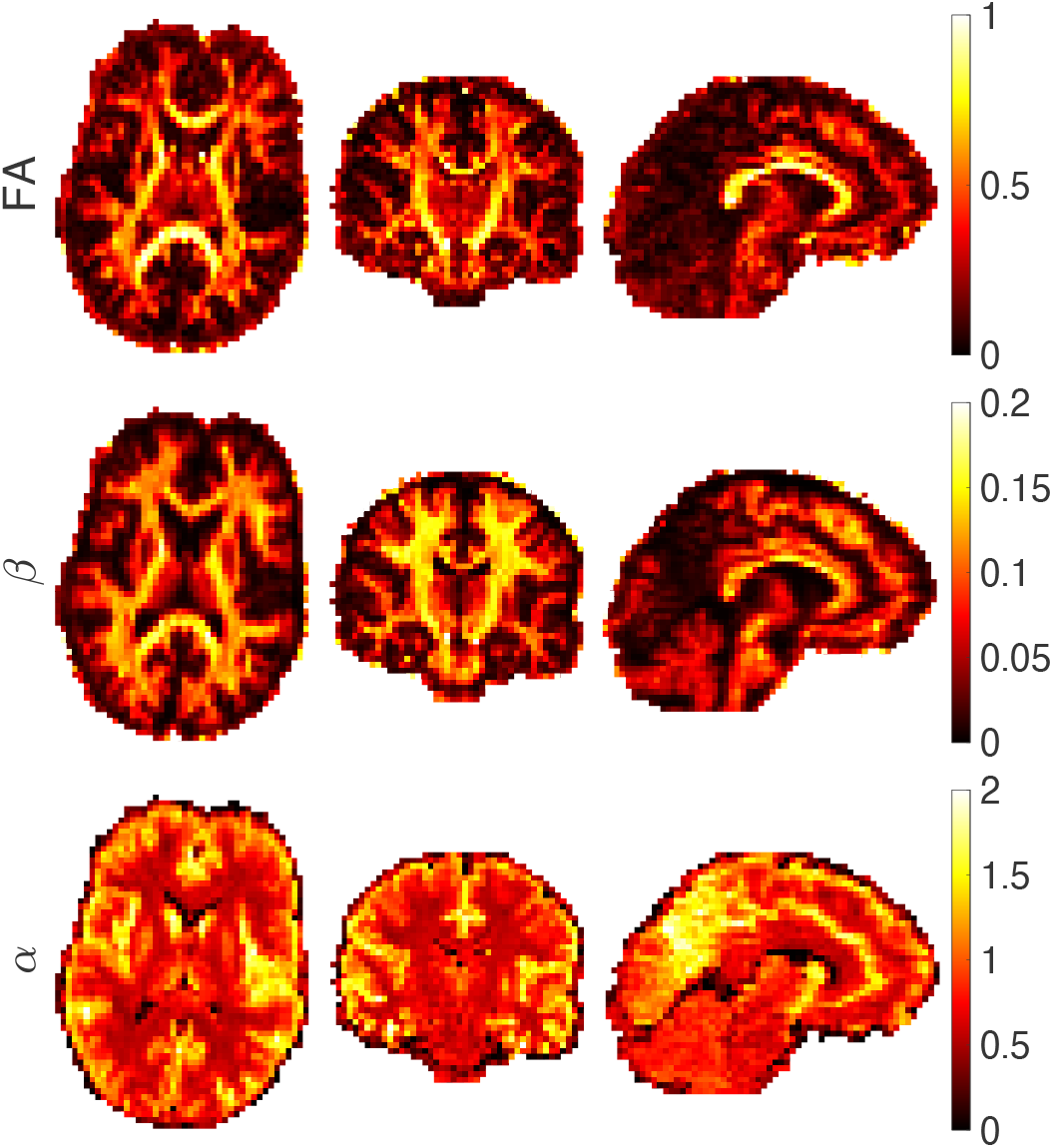
Estimated FA, parameter *β* and *α* of the power-law fit (*S/S*_0_ = *βb^−α^*) from axial, coronal, and sagittal views of the brain image.

## 5. Discussion

The SANDI model (Palombo et al., 2020) extends existing multi-compartment models that only consider two pools of water in brain tissue (Jespersen et al., 2007; Zhang et al., 2012; Fieremans et al., 2011; Kaden et al., 2016; Novikov et al., 2018a, 2019; Alexander et al., 2019). Palombo et al. (Palombo et al., 2018a) suggested that the failure of the stick model in gray matter (McKinnon et al., 2017; Veraart et al., 2019) can be due to the abundance of cell bodies (namely soma). In previous works, the contribution from soma was considered as part of the extracellular space (Jespersen et al., 2007; Jespersen, 2012) because the exchange between the restricted water in soma and the hindered water in the extracellular space was assumed to be fast. However, recent studies (Yang et al., 2018a) suggest that the pre-exchange time of intracellular water in neurons and astrocytes is ~500 *ms*. These findings prompt the conclusion that for diffusion times much smaller than 500 *ms* (e.g. ~10-20 ms) the exchange between intra and extra-cellular water may be negligible, supporting the idea that the signal from spins restricted in soma may be non-negligible.

The typical size of soma ranges from a few microns (for microglia and glia) to a few tens of microns (for big neurons) (Zhao et al., 2006; Díaz-Cintra et al., 2004). In this study we found that based on the power-law fit to the signal in voxels containing white matter, the sphere radius is larger than 7*μm* (Fig. .6). While at first glance, one might not expect spherical compartments of this size in white matter *in vivo*, there are studies that report the size of astrocytes (Savtchenko et al., 2018; Di Benedetto et al., 2016; Zhang et al., 2016; Papageorgiou et al., 2011) and oligodendrocytes (Fannon et al., 2015; Mohamed et al., 2020) and confirm that having effective soma radius > 7*μm* is possible (data also publicly available on Neuromorpho.org). This suggests the presence of glial cells in the central nervous system whose soma can have such a size. Although in healthy conditions glial volume fraction in white matter has been estimated to be low (5% 20% (Veraart et al., 2020; Coelho et al., 2018; Duval et al., 2016)), the presence of large glial cells may become more relevant in pathological conditions, where glia may migrate and react undergoing hypertrophy of the whole cellular structure (Ligneul et al., 2019; Genovese et al., 2020).

In this work, we studied the minimal sphere signal fraction and radius that can be detected from diffusion MRI of water inside spheres. Our finding is in agreement with the results reported by Dhital et al. (Dhital et al., 2018) (2% of the unweighted signal for moderate diffusion times using Prisma scanner) and Tax et al. (Tax et al., 2020) (isotropic signal fraction of 9.7% for the apparent diffusivity of 0.12*μm*^2^/*ms*). It should be noted that the protocol in this study is not specifically optimized for size estimation. For example, intentionally varying the frequency spectra (e.g. to include high frequency components), might result in better sensitivity to smaller pore-sizes (Molendowska et al., 2020).

The estimation of stick signal fraction in (Lampinen et al., 2019) was challenging while in (Lampinen et al., 2020) Lampinen et al. provided reliable estimates of the two-compartment model parameters because the sequence was optimized in the latter. Comparing the estimation of sphere signal fraction in this work with the stick signal fraction in (Lampinen et al., 2019, 2020) we conclude that to estimate the parameters accurately, the acquisition should be optimized toward the estimation of the model parameters.

Here we used a combination of linear, planar, and spherical tensor encoding to ameliorate the degeneracy problem that exists in the fitting of multi-compartment models. Nilsson et al. (Nilsson et al., 2017) reported that for the estimation of diameter in complete orientation dispersion (which we effect by powder-averaging the signal), from an SNR perspective it is advantageous to use oscillating gradient spin echo (OGSE) compared to standard SDE. The benefits of double diffusion encoding (DDE) for size estimation have been presented in other studies (Benjamini et al., 2014; Katz and Nevo, 2014; Vincent et al., 2020). Our results suggest using multiple waveforms provides the best estimates.

In the case of cylinder diameter estimation, the resolution limit is determined by the amount of attenuation due to radial diffusion. This attenuation is estimated by the integral of the gradient squared and can be maximized by either a fat-pulse SDE or a rectangular oscillating pulse. However, when the long axis of the cylinder is not perpendicular to the direction of the applied gradient, the high b-values should be avoided because of the signal attenuation and decrease in SNR. To improve the SNR and the resolution limit for cylinder diameter estimation, waveforms should have more oscillations and hence lower b-values (Nilsson et al., 2017). Here, we are targeting the sphere diameter, and therefore OGSE or SDE can be both useful.

In the estimation of spherical pore size from the diffusion-weighted signal, different confounding factors have to be considered. One of the challenges is that for small sphere radii (< 3 micron), the fitting landscape is flat and there is a negligible change in the signal for small sphere sizes (Fig. .3). Noise is another confounding factor that affects the estimation of parameters in both model-based and signal representation based techniques. Parameters obtained from multi-compartment models, (the stick + ball + sphere model in this paper) applied to noisy data are biased because of the effect of noise. Three different noise scenarios were simulated here: Gaussian noise, Rician noise, and corrected Rician noise. If the data were corrupted with purely Gaussian noise, then this could be removed to some extent through the act of computing the orientationally-averaged signal. However, as we invariably use the magnitude-reconstructed data, the noise has a Rician distribution, which presents a more challenging scenario because averaging does not remove the bias. This effect is more pronounced when there is a small contribution from the spherical compartment.

We wish to stress that the challenges we identified are mainly relevant to WM. The SANDI model was developed mainly for soma imaging in GM. Nothing in our results suggests that SANDI is unreliable in GM, and will indeed, benefit from the multi-waveform and frequency-domain approach presented here. The challenge in the gray matter is to determine whether the deviation from the ‘stick’ model comes from the soma compartment, exchange between compartments (Jelescu and Novikov, 2020) or both. We note that the extracellular signal fraction obtained from the fit in the gray matter areas is around 0.45 which is higher than expected. This discrepancy can be explained by three factors: first, *T*_2_ relaxation is not explicitly accounted for in our model, and second, we consider the same proton density in both intra-axonal and extra-axonal spaces. And also non-negligible partial volume with CSF, particularly problematic for cortical GM at 3 *mm*^3^ resolution. The model presented here is only sensitive to relative signal amplitudes while differences in *T*_2_ relaxation can impose different weights to the amplitudes of the pools (Szafer et al., 1995; Callaghan, 1995). The specific assignment of nerve water *T*_2_ components by simultaneously considering compartmental diffusion and transverse relaxation properties was already studied by Peled et al. (Peled et al., 1999) in myelinated nerves of the frog sciatic nerve tissue. More and more evidence is given for the *T*_2_ relaxation time constants of intra- and extra-axonal water to be different from each other in case of slow exchange between the intra-axonal and extra-axonal pools (Peled et al., 1999; Dortch et al., 2010). In the case of fast exchange (Szafer et al., 1995), the signal loss in all the pools would be weighted in the same manner. Although it is relatively easy to incorporate relaxation into the model and fit the experimental data with additional parameters (relaxation rates), such a model results in an unstable fit with the current protocol. To incorporate additional parameters (i.e., relaxation rates) in the model we need to obtain additional information in our experiment. For example, repeating the acquisition at different echo times as suggested by Lampinen et al. (Lampinen et al., 2020).

We highlight six limitations of the current study in the estimation of spherical compartment signal fraction and size. First, we assume that the intra-axonal water comes from straight axons, which is not the case in most of the white matter voxels (Nilsson et al., 2012; Nilsson and Alexander, 2012; Jeurissen et al., 2013; Lee et al., 2020; Özarslan et al., 2018). Second, the extra-cellular component is modelled as a ball with isotropic diffusion. This assumption is not valid when there are coherently-oriented fibers (such as in the midline of the corpus callosum), where diffusion in the extra-axonal space can have a high anisotropy (Özarslan et al., 2011). Third, at low frequencies, the time-dependency of diffusivity in the extracellular space can dominate over the time-dependency in the intra-axonal space (Burcaw et al., 2015; Nilsson et al., 2017). Fourth, it is assumed that the exchange between water environments is negligible. This assumption might be valid since previous studies have shown the exchange times in the white matter are of the order of seconds or longer (Nilsson et al., 2013) which is much larger than the time-scales that the effects of restricted diffusion can be observed. Fifth, in our tissue model, we have neglected the distribution of restricted dimensions (e.g. range of soma sizes, axon diameters). However, adding extra parameters to the model to account for this will make the fitting unstable. Six, the effect of *T*_2_ relaxation is not considered in our model which may result in bias in the estimation of the other parameters of the model. In addition, the lack of cerebrospinal fluid (CSF) component in the model is another limitation of this work.

### Future directions

The fitting method used in this work is a nonlinear least-square fit that can be replaced with new deep learning approaches to improve the quality of fit (Gibbons et al., 2019). From the acquisition perspective, the protocol that is used in this study is not optimized for the estimation of small sizes. Using a range of frequency spectra will help (Molendowska et al., 2020). The protocol, used in this work, imposes a long acquisition time which can be minimized by optimizing the directions as well as the number of shells. In this paper, simple arithmetic averaging is used for powder averaging which can be replaced with some other techniques such as weighted averaging (Knutsson et al., 1999; Szczepankiewicz et al., 2017; Afzali et al., 2020b) to obtain a better orientation-invariant signal that improves the parameter estimates.

## 6. Conclusion

In this work, we have demonstrated key challenges and limitations in estimating spherical pore size non-invasively in the human brain from diffusion MRI. Our simulations show the effect of Rician bias on the estimation of pore size and identified the lower bound limit of the sphere signal fraction and size that can be detected from the diffusion-weighted signal from both an SNR and empirical perspective. We showed that for small signal fraction of soma, i.e. < 10%, this is a problem. However, we know from detailed microscopy of brain cortex (Beul and Hilgetag, 2019; Collins et al., 2010), that in GM the soma signal fraction is on average > ~20%. Therefore, reliable estimation of soma properties in GM is possible, while in WM it presents several challenges. The flat landscape of the fitting was also investigated. Using the ultra-strong gradients of the Connectom scanner, the diffusion signal in the white matter can be made sensitive to the axon diameter, and therefore the three-compartment model of stick + ball + sphere changes to cylinder + ball + sphere which has two time-dependencies, one for the diffusivity in the sphere and the other one for the diffusivity in the cylinder. Disentangling these two time-dependencies using only one sequence parameter (i.e., changing the frequency content of the encoding waveform) in the acquisition is challenging. Studying all these challenges prevents misinterpretation of the biased estimated parameters as the potential biomarkers in clinical studies.

## Acknowledgements

The data were acquired at the UK National Facility for In Vivo MR Imaging of Human Tissue Microstructure funded by the EPSRC (grant EP/M029778/1), and The Wolfson Foundation. MA and DKJ are supported by a Wellcome Trust Investigator Award (096646/Z/11/Z) and a Wellcome Trust Strategic Award (104943/Z/14/Z). MN was supported by grants from Swedish Research Council (2016-03443) and Random Walk Imaging AB (grant no. MN15). The authors would like to thank Filip Szczepankiewicz for providing the pulse sequences for b-tensor encoding. We thank Lars Mueller for setting up the protocol for b-tensor encoding. MP is supported by UK EPSRC EP/N018702/1 and UKRI Future Leaders Fellowship MR/T020296/1.

## A. Supplementary Material

Fig. S1 shows the estimated parameters of the stick + ball + sphere model on axial, coronal, and sagittal views of the *in vivo* brain image without applying the Gaussian kernel. In Fig. S2, in addition to the parameters of the model we also estimate the standard deviation of the noise as it is explained in Section 3.2.

**Figure S1:**
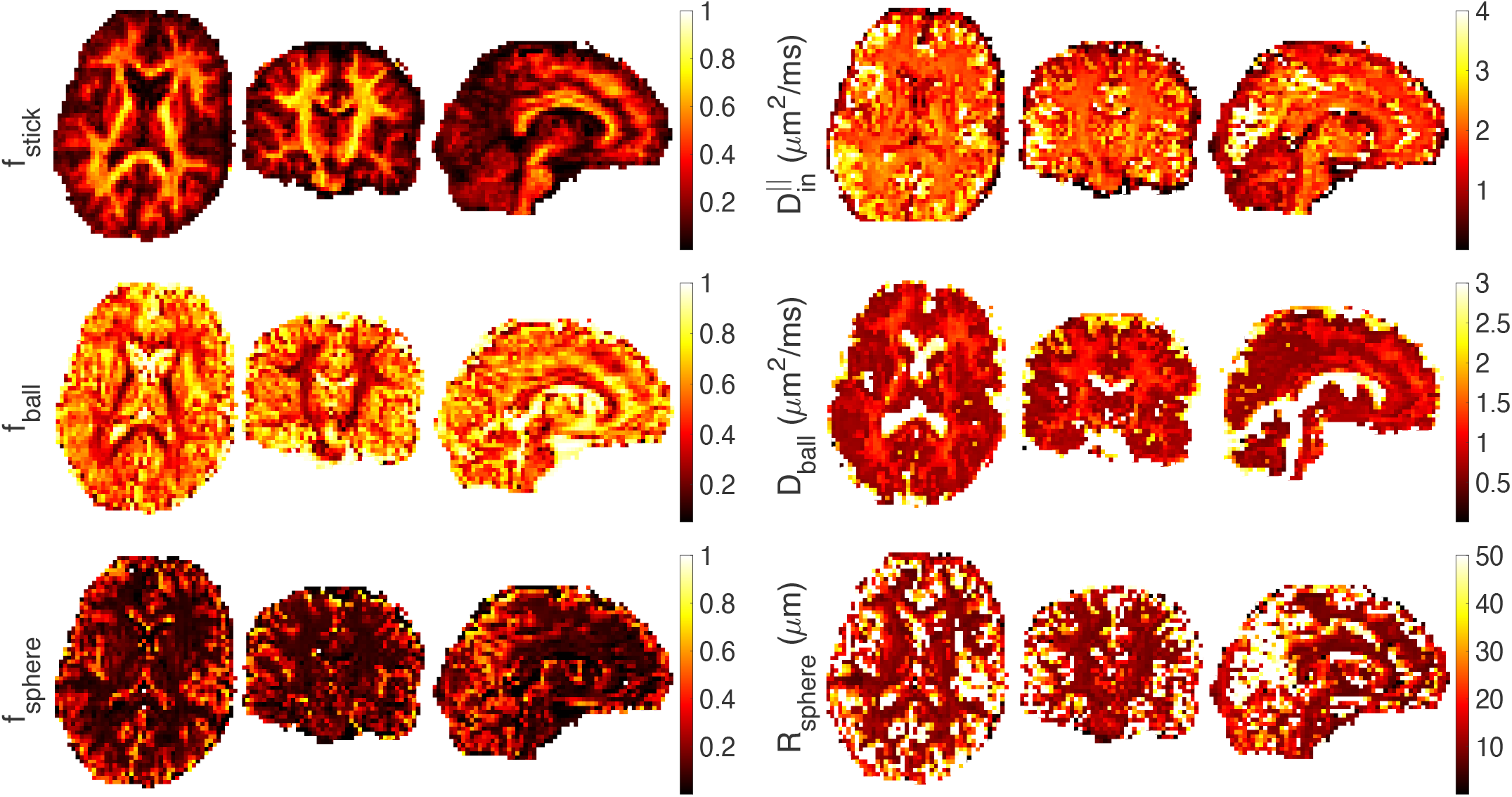
Estimated stick (*f*_stick_), ball (*f*_ball_), and sphere (*f*_sphere_) signal fractions, intra-axonal parallel diffusivity 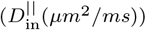, extra-cellular diffusivity (*D*_ball_(*μm*^2^/*ms*)), and sphere radius (*R*_sphere_(*μm*)) on axial, coronal, and sagittal views of the *in vivo* brain image.

**Figure S2:**
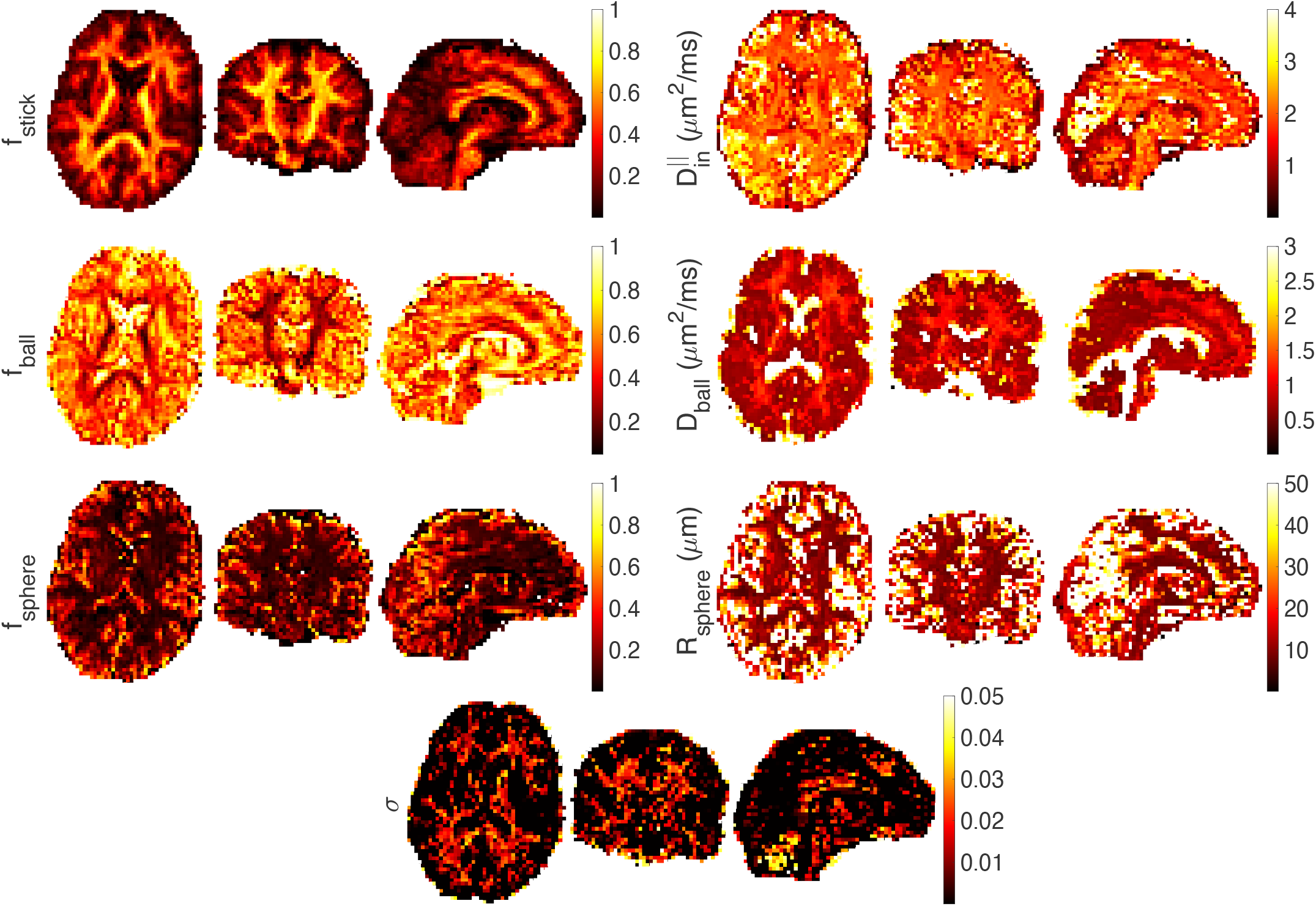
Estimated stick (*f*_stick_), ball (*f*_ball_), and sphere (*f*_sphere_) signal fractions, intra-axonal parallel diffusivity 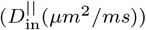, extra-cellular diffusivity (*D*_ball_(*μm*^2^/*ms*)), sphere radius (*R*_sphere_(*μm*)), and standard deviation of the noise (*σ*) on axial, coronal, and sagittal views of the *in vivo* brain image.

## References

Afzali, M., Aja-Fernández, S., Jones, D.K.. Direction-averaged diffusion-weighted MRI signal using different axisymmetric B-tensor encoding schemes. Magnetic Resonance in Medicine 2020a;.

Afzali, M., Knutsson, H., Özarslan, E., Jones, D.K.. Computing the orientational-average of diffusion-weighted mri signals: A comparison of different techniques. bioRxiv 2020b;.

Afzali, M., Pieciak, T., Newman, S., Garifallidis, E., Özarslan, E., Cheng, H., Jones, D.K.. The sensitivity of diffusion MRI to microstructural properties and experimental factors. Journal of Neuroscience Methods 2020c;:108951.

Aja-Fernández, S., Vegas-Sánchez-Ferrero, G.. Statistical analysis of noise in MRI. Switzerland: Springer International Publishing 2016;.

Alexander, D.C., Dyrby, T.B., Nilsson, M., Zhang, H.. Imaging brain microstructure with diffusion MRI: practicality and applications. NMR in Biomedicine 2019;32(4):e3841.

Andersson, J.L., Sotiropoulos, S.N.. An integrated approach to correction for off-resonance effects and subject movement in diffusion mr imaging. Neuroimage 2016;125:1063–1078.

Andrae, R., Schulze-Hartung, T., Melchior, P.. Dos and don’ts of reduced chi-squared. arXiv preprint arXiv:10123754 2010;.

Assaf, Y., Blumenfeld-Katzir, T., Yovel, Y., Basser, P.. Axcaliber: a method for measuring axon diameter distribution from diffusion MRI. Magnetic Resonance in Medicine: An Official Journal of the International Society for Magnetic Resonance in Medicine 2008;59(6):1347–1354.

Behrens, T.E., Woolrich, M.W., Jenkinson, M., Johansen-Berg, H., Nunes, R.G., Clare, S., Matthews, P.M., Brady, J.M., Smith, S.M.. Characterization and propagation of uncertainty in diffusion-weighted mr imaging. Magnetic Resonance in Medicine: An Official Journal of the International Society for Magnetic Resonance in Medicine 2003;50(5):1077–1088.

Benjamini, D., Komlosh, M.E., Basser, P.J., Nevo, U.. Nonparametric pore size distribution using d-pfg: comparison to s-pfg and migration to MRI. Journal of Magnetic Resonance 2014;246:36–45.

Beul, S.F., Hilgetag, C.C.. Neuron density fundamentally relates to architecture and connectivity of the primate cerebral cortex. NeuroImage 2019;189:777–792.

Burcaw, L.M., Fieremans, E., Novikov, D.S.. Mesoscopic structure of neuronal tracts from time-dependent diffusion. NeuroImage 2015;114:18–37.

Callaghan, P., Jolley, K., Lelievre, J.. Diffusion of water in the endosperm tissue of wheat grains as studied by pulsed field gradient nuclear magnetic resonance. Biophysical journal 1979;28(1):133–141.

Callaghan, P.T.. Pulsed-gradient spin-echo NMR for planar, cylindrical, and spherical pores under conditions of wall relaxation. Journal of magnetic resonance, Series A 1995;113(1):53–59.

Callaghan, P.T.. Translational dynamics and magnetic resonance: Principles of pulsed gradient spin echo NMR. New York: Oxford University Press, 2011.

Coelho, S., Pozo, J.M., Costantini, M., Highley, J.R., Mozumder, M., Simpson, J.E., Ince, P.G., Frangi, A.F.. Local volume fraction distributions of axons, astrocytes, and myelin in deep subcortical white matter. Neuroimage 2018;179:275–287.

Coelho, S., Pozo, J.M., Jespersen, S.N., Jones, D.K., Frangi, A.F.. Resolving degeneracy in diffusion MRI biophysical model parameter estimation using double diffusion encoding. Magnetic resonance in medicine 2019;82(1):395–410.

Collins, C.E., Airey, D.C., Young, N.A., Leitch, D.B., Kaas, J.H.. Neuron densities vary across and within cortical areas in primates. Proceedings of the National Academy of Sciences 2010;107(36):15927–15932.

Cory, D., Garroway, A., Miller, J.. Applications of spin transport as a probe of local geometry. In: Abstracts of Papers of the American Chemical Society. AMER CHEMICAL SOC 1155 16TH ST, NW, WASHINGTON, DC 20036; volume 199; 1990. p. 105–POLY.

Danos, P., Baumann, B., Krämer, A., Bernstein, H.G., Stauch, R., Krell, D., Falkai, P., Bogerts, B.. Volumes of association thalamic nuclei in schizophrenia: a postmortem study. Schizophrenia Research 2003;60(2-3):141–155.

Dhital, B., Kellner, E., Kiselev, V.G., Reisert, M.. The absence of restricted water pool in brain white matter. Neuroimage 2018;182:398–406.

Dhital, B., Reisert, M., Kellner, E., Kiselev, V.G.. Intra-axonal diffusivity in brain white matter. NeuroImage 2019;189:543–550.

Di Benedetto, B., Malik, V.A., Begum, S., Jablonowski, L., Gómez-González, G.B., Neumann, I.D., Rupprecht, R.. Fluoxetine requires the endfeet protein aquaporin-4 to enhance plasticity of astrocyte processes. Frontiers in cellular neuroscience 2016;10:8.

Díaz-Cintra, S., Yong, A., Aguilar, A., Bi, X., Lynch, G., Ribak, C.E.. Ultrastructural analysis of hippocampal pyramidal neurons from apolipoprotein e-deficient mice treated with a cathepsin inhibitor. Journal of neurocytology 2004;33(1):37–48.

Dortch, R.D., Apker, G.A., Valentine, W.M., Lai, B., Does, M.D.. Compartment-specific enhancement of white matter and nerve ex vivo using chromium. Magnetic resonance in medicine 2010;64(3):688–697.

Duval, T., Stikov, N., Cohen-Adad, J.. Modeling white matter microstructure. Functional neurology 2016;31(4):217.

Edén, M.. Computer simulations in solid-state NMR. iii. powder averaging. Concepts in Magnetic Resonance Part A: An Educational Journal 2003;18(1):24–55.

Eriksson, S., Lasič, S., Nilsson, M., Westin, C.F., Topgaard, D.. NMR diffusion-encoding with axial symmetry and variable anisotropy: Distinguishing between prolate and oblate microscopic diffusion tensors with unknown orientation distribution. The Journal of chemical physics 2015;142(10):104201.

Fannon, J., Tarmier, W., Fulton, D.. Neuronal activity and ampa-type glutamate receptor activation regulates the morphological development of oligodendrocyte precursor cells. Glia 2015;63(6):1021–1035.

Fieremans, E., Jensen, J.H., Helpern, J.A.. White matter characterization with diffusional kurtosis imaging. Neuroimage 2011;58(1):177–188.

Fieremans, E., Veraart, J., Benjamin, A., Filip, S., Nilsson, M., Novikov, D.. Effect of combining linear with spherical tensor encoding on estimating brain microstructural parameters. Proceedings of the ISMRM, Paris 2018;.

Genovese, G., Palombo, M., Santin, M.D., Valette, J., Ligneul, C., Aigrot, M.S., Abdoulkader, N., Langui, D., Millecamps, A., Evercooren, A.B.V., et al. Inflammation-driven glial alterations in the cuprizone mouse model probed with diffusion-weighted magnetic resonance spectroscopy at 11.7 t. arXiv preprint arXiv:200708400 2020;.

Gibbons, E.K., Hodgson, K.K., Chaudhari, A.S., Richards, L.G., Majersik, J.J., Adluru, G., DiBella, E.V.. Simultaneous noddi and gfa parameter map generation from subsampled q-space imaging using deep learning. Magnetic resonance in medicine 2019;81(4):2399–2411.

Gyori, N., Clark, C., Dragonu, I., Alexander, D., Kaden, E.. In-vivo neural soma imaging using b-tensor encoding and deep learning. In: Proceedings of the 27th Annual Meeting of ISMRM, Montreal, Canada. 2019..

Henriques, R.N., Jespersen, S.N., Shemesh, N.. Microscopic anisotropy misestimation in spherical-mean single diffusion encoding MRI. Magnetic resonance in medicine 2019;81(5):3245–3261.

Ianuş, A., Drobnjak, I., Alexander, D.C.. Model-based estimation of microscopic anisotropy using diffusion MRI: a simulation study. NMR in Biomedicine 2016;29(5):672–685.

Jelescu, I., Novikov, D.. Water exchange time between gray matter compartments in vivo. In: Proceedings of the 29th Annual Meeting of ISMRM. 2020..

Jelescu, I., Veraart, J., Fieremans, E., Novikov, D.. Degeneracy in model parameter estimation for multi-compartmental diffusion in neuronal tissue. NMR in Biomedicine 2016;29(1):33–47.

Jespersen, S.N.. Equivalence of double and single wave vector diffusion contrast at low diffusion weighting. NMR in Biomedicine 2012;25(6):813–818.

Jespersen, S.N., Kroenke, C.D., Østergaard, L., Ackerman, J.J., Yablonskiy, D.A.. Modeling dendrite density from magnetic resonance diffusion measurements. Neuroimage 2007;34(4):1473–1486.

Jespersen, S.N., Lundell, H., Sønderby, C.K., Dyrby, T.B.. Orientationally invariant metrics of apparent compartment eccentricity from double pulsed field gradient diffusion experiments. NMR in Biomedicine 2013;26(12):1647–1662.

Jespersen, S.N., Olesen, J.L., Ianuş, A., Shemesh, N.. Effects of nongaussian diffusion on isotropic diffusion measurements: an ex-vivo microimaging and simulation study. Journal of Magnetic Resonance 2019;300:84–94.

Jeurissen, B., Leemans, A., Tournier, J.D., Jones, D.K., Sijbers, J.. Investigating the prevalence of complex fiber configurations in white matter tissue with diffusion magnetic resonance imaging. Human brain mapping 2013;34(11):2747–2766.

Jones, D.K., Basser, P.J.. Squashing peanuts and smashing pumpkins”: how noise distorts diffusion-weighted mr data. Magnetic Resonance in Medicine: An Official Journal of the International Society for Magnetic Resonance in Medicine 2004;52(5):979–993.

Jones, D.K., Knösche, T.R., Turner, R.. White matter integrity, fiber count, and other fallacies: the do’s and don’ts of diffusion MRI. Neuroimage 2013;73:239–254.

Kaden, E., Kruggel, F., Alexander, D.C.. Quantitative mapping of the per-axon diffusion coefficients in brain white matter. Magnetic resonance in medicine 2016;75(4):1752–1763.

Katz, Y., Nevo, U.. Quantification of pore size distribution using diffusion NMR: experimental design and physical insights. The Journal of Chemical Physics 2014;140(16):164201.

Kellner, E., Dhital, B., Kiselev, V.G., Reisert, M.. Gibbs-ringing artifact removal based on local subvoxel-shifts. Magnetic resonance in medicine 2016;76(5):1574–1581.

Knutsson, H.. Towards optimal sampling in diffusion MRI. In: International Conference on Medical Image Computing and Computer-Assisted Intervention. Springer; 2018. p. 3–18.

Knutsson, H., Andersson, M., Wiklund, J.. Advanced filter design. SCIA; 1999..

Koay, C.G., Özarslan, E., Basser, P.J.. A signal transformational framework for breaking the noise floor and its applications in MRI. Journal of magnetic resonance 2009;197(2):108–119.

Lampinen, B., Szczepankiewicz, F., Mårtensson, J., van Westen, D., Hansson, O., Westin, C.F., Nilsson, M.. Towards unconstrained compartment modeling in white matter using diffusion-relaxation MRI with tensor-valued diffusion encoding. Magnetic Resonance in Medicine 2020;84(3):1605–1623.

Lampinen, B., Szczepankiewicz, F., Mårtensson, J., van Westen, D., Sundgren, P.C., Nilsson, M.. Neurite density imaging versus imaging of microscopic anisotropy in diffusion MRI: a model comparison using spherical tensor encoding. Neuroimage 2017;147:517–531.

Lampinen, B., Szczepankiewicz, F., Novén, M., van Westen, D., Hansson, O., Englund, E., Mårtensson, J., Westin, C.F., Nilsson, M.. Searching for the neurite density with diffusion MRI: challenges for biophysical modeling. Human brain mapping 2019;40(8):2529–2545.

Lasič, S., Szczepankiewicz, F., Eriksson, S., Nilsson, M., Topgaard, D.. Microanisotropy imaging: quantification of microscopic diffusion anisotropy and orientational order parameter by diffusion MRI with magic-angle spinning of the q-vector. Frontiers in Physics 2014;2:11.

Lee, H.H., Papaioannou, A., Kim, S.L., Novikov, D.S., Fieremans, E.. A time-dependent diffusion MRI signature of axon caliber variations and beading. Communications biology 2020;3(1):1–13.

Ligneul, C., Palombo, M., Hernández-Garzón, E., Carrillo-de Sauvage, M.A., Flament, J., Hantraye, P., Brouillet, E., Bonvento, G., Escartin, C., Valette, J.. Diffusion-weighted magnetic resonance spectroscopy enables cell-specific monitoring of astrocyte reactivity in vivo. NeuroImage 2019;191:457–469.

Lundell, H., Nilsson, M., Dyrby, T., Parker, G., Cristinacce, P.H., Zhou, F.L., Topgaard, D., Lasič, S.. Multidimensional diffusion MRI with spectrally modulated gradients reveals unprecedented microstructural detail. Scientific reports 2019;9(1):9026.

McKinnon, E.T., Jensen, J.H., Glenn, G.R., Helpern, J.A.. Dependence on b-value of the direction-averaged diffusion-weighted imaging signal in brain. Magnetic resonance imaging 2017;36:121–127.

Mitra, P.P., Sen, P.N., Schwartz, L.M., Le Doussal, P.. Diffusion propagator as a probe of the structure of porous media. Physical review letters 1992;68(24):3555.

Mohamed, E., Paisley, C.E., Meyer, L.C., Bigbee, J.W., Sato-Bigbee, C.. Endogenous opioid peptides and brain development: Endomorphin-1 and nociceptin play a sex-specific role in the control of oligodendrocyte maturation and brain myelination. Glia 2020;68(7):1513–1530.

Molendowska, M., Drakesmith, M., Jones, D., Tax, C.. Varying the frequency-content of high b-value spherical diffusion encoding improves the characterisation of isotropic restricted compartment. In: Proceedings of the 29th Annual Meeting of ISMRM. 2020..

Mori, S., Wakana, S., Van Zijl, P.C., Nagae-Poetscher, L.. MRI atlas of human white matter. Elsevier, 2005.

Murday, J., Cotts, R.M.. Self-diffusion coefficient of liquid lithium. The Journal of Chemical Physics 1968;48(11):4938–4945.

Nilsson, M., Alexander, D.. Investigating tissue microstructure using diffusion MRI: How does the resolution limit of the axon diameter relate to the maximal gradient strength. In: Proc Intl Soc Mag Reson Med. volume 20; 2012. p. 3567.

Nilsson, M., Lasič, S., Drobnjak, I., Topgaard, D., Westin, C.F.. Resolution limit of cylinder diameter estimation by diffusion MRI: The impact of gradient waveform and orientation dispersion. NMR in Biomedicine 2017;30(7):e3711.

Nilsson, M., Lätt, J., Ståhlberg, F., van Westen, D., Hagslätt, H.. The importance of axonal undulation in diffusion mr measurements: a monte carlo simulation study. NMR in Biomedicine 2012;25(5):795–805.

Nilsson, M., Lätt, J., van Westen, D., Brockstedt, S., Lasič, S., Ståhlberg, F., Topgaard, D.. Noninvasive mapping of water diffusional exchange in the human brain using filter-exchange imaging. Magnetic resonance in medicine 2013;69(6):1572–1580.

Novikov, D., Veraart, J., Jelescu, I., Fieremans, E.. Rotationally-invariant mapping of scalar and orientational metrics of neuronal microstructure with diffusion MRI. NeuroImage 2018a;174:518–538.

Novikov, D.S., Fieremans, E., Jespersen, S.N., Kiselev, V.G.. Quantifying brain microstructure with diffusion MRI: Theory and parameter estimation. NMR in Biomedicine 2019;32(4):e3998.

Novikov, D.S., Kiselev, V.G., Jespersen, S.N.. On modeling. Magnetic resonance in medicine 2018b;79(6):3172–3193.

Özarslan, E., Shemesh, N., Basser, P.J.. A general framework to quantify the effect of restricted diffusion on the NMR signal with applications to double pulsed field gradient NMR experiments. The Journal of chemical physics 2009;130(10):104702.

Özarslan, E., Shemesh, N., Koay, C.G., Cohen, Y., Basser, P.J.. Nuclear magnetic resonance characterization of general compartment size distributions. New journal of physics 2011;13(1):015010.

Özarslan, E., Yolcu, C., Herberthson, M., Knutsson, H., Westin, C.F.. Influence of the size and curvedness of neural projections on the orientationally averaged diffusion mr signal. Frontiers in physics 2018;6:17.

Palombo, M., Ianus, A., Guerreri, M., Nunes, D., Alexander, D.C., Shemesh, N., Zhang, H.. SANDI: a compartment-based model for non-invasive apparent soma and neurite imaging by diffusion MRI. NeuroImage 2020;:116835.

Palombo, M., Shemesh, N., Ianus, A., Alexander, D., Zhang, H.. Abundance of cell bodies can explain the stick models failure in grey matter at high bvalue. ISMRM (International Society for Magnetic Resonance in Medicine); 2018a..

Palombo, M., Shemesh, N., Ianus, A., Alexander, D.C., Zhang, H.. A compartment based model for non-invasive cell body imaging by diffusion MRI. In: Proceedings of the International Society for Magnetic Resonance in Medicine. volume 27; 2018b. p. 580.

Panagiotaki, E., Schneider, T., Siow, B., Hall, M.G., Lythgoe, M.F., Alexander, D.C.. Compartment models of the diffusion mr signal in brain white matter: a taxonomy and comparison. Neuroimage 2012;59(3):2241–2254.

Papageorgiou, I.E., Gabriel, S., Fetani, A.F., Kann, O., Heinemann, U.. Redistribution of astrocytic glutamine synthetase in the hippocampus of chronic epileptic rats. Glia 2011;59(11):1706–1718.

Peled, S., Cory, D.G., Raymond, S.A., Kirschner, D.A., Jolesz, F.A.. Water diffusion, t2, and compartmentation in frog sciatic nerve. Magnetic Resonance in Medicine: An Official Journal of the International Society for Magnetic Resonance in Medicine 1999;42(5):911–918.

Pieciak, T., Aja-Fernández, S., Vegas-Sánchez-Ferrero, G.. Non-stationary rician noise estimation in parallel mri using a single image: a variance-stabilizing approach. IEEE Transactions on Pattern Analysis and Machine Intelligence 2016a;39(10):2015–2029.

Pieciak, T., Rabanillo-Viloria, I., Aja-Fernández, S.. Bias correction for non-stationary noise filtering in mri. In: 2018 IEEE 15th International Symposium on Biomedical Imaging (ISBI 2018). IEEE; 2018. p. 307–310.

Pieciak, T., Vegas-Sánchez-Ferrero, G., Aja-Fernández, S.. Variance stabilization of noncentral-chi data: Application to noise estimation in mri. In: 2016 IEEE 13th International Symposium on Biomedical Imaging (ISBI). IEEE; 2016b. p. 1376–1379.

Reisert, M., Kiselev, V.G., Dhital, B.. A unique analytical solution of the white matter standard model using linear and planar encodings. Magnetic resonance in medicine 2019;81(6):3819–3825.

Rudrapatna, S., Parker, G., Roberts, J., Jones, D.. Can we correct for interactions between subject motion and gradient-nonlinearity in diffusion MRI. In: Proc. Int. Soc. Mag. Reson. Med. volume 1206; 2018..

Rudrapatna, U., Parker, G.D., Jamie, R., Jones, D.K.. A comparative study of gradient nonlinearity correction strategies for processing diffusion data obtained with ultra-strong gradient MRI scanners. Magnetic Resonance in Medicine 2020;.

Savtchenko, L.P., Bard, L., Jensen, T.P., Reynolds, J.P., Kraev, I., Medvedev, N., Stewart, M.G., Henneberger, C., Rusakov, D.A.. Disentangling astroglial physiology with a realistic cell model in silico. Nature communications 2018;9(1):1–15.

Sjölund, J., Szczepankiewicz, F., Nilsson, M., Topgaard, D., Westin, C.F., Knutsson, H.. Constrained optimization of gradient waveforms for generalized diffusion encoding. Journal of Magnetic Resonance 2015;261:157–168.

Stanisz, G., Wright, G., Henkelman, R., Szafer, A.. An analytical model of restricted diffusion in bovine optic nerve. Magnetic Resonance in Medicine 1997;37(1):103–111.

Stejskal, E.O., Tanner, J.E.. Spin diffusion measurements: spin echoes in the presence of a time-dependent field gradient. The journal of chemical physics 1965;42(1):288–292.

Stepišnik, J.. Time-dependent self-diffusion by NMR spin-echo. Physica B: Condensed Matter 1993;183(4):343–350.

Szafer, A., Zhong, J., Gore, J.C.. Theoretical model for water diffusion in tissues. Magnetic resonance in medicine 1995;33(5):697–712.

Szczepankiewicz, F., Westin, C.F., Knutsson, H.. A measurement weighting scheme for optimal powder average estimation. In: Proc Intl Soc Mag Reson Med. volume 26; 2017. p. 3345.

Szczepankiewicz, F., Westin, C.F., Nilsson, M.. Maxwell-compensated design of asymmetric gradient waveforms for tensor-valued diffusion encoding. Magnetic resonance in medicine 2019;82(4):1424–1437.

Tax, C.M., Szczepankiewicz, F., Nilsson, M., Jones, D.K.. The dot-compartment revealed? diffusion MRI with ultra-strong gradients and spherical tensor encoding in the living human brain. NeuroImage 2020;210:116534.

Topgaard, D.. Multidimensional diffusion MRI. Journal of Magnetic Resonance 2017;275:98–113.

Tziortzi, A.C., Searle, G.E., Tzimopoulou, S., Salinas, C., Beaver, J.D., Jenkinson, M., Laruelle, M., Rabiner, E.A., Gunn, R.N.. Imaging dopamine receptors in humans with [11c]-(+)-phno: dissection of d3 signal and anatomy. Neuroimage 2011;54(1):264–277.

Vangelderen, P., DesPres, D., Vanzijl, P., Moonen, C.. Evaluation of restricted diffusion in cylinders. phosphocreatine in rabbit leg muscle. Journal of Magnetic Resonance, Series B 1994;103(3):255–260.

Veraart, J., Fieremans, E., Novikov, D.S.. On the scaling behavior of water diffusion in human brain white matter. NeuroImage 2019;185:379–387.

Veraart, J., Fieremans, E., Rudrapatna, U., Jones, D., Novikov, D.. Biophysical modeling of the gray matter: does the “stick” model hold. In: Proceedings of the 27th Annual Meeting of ISMRM, Paris, France. 2018..

Veraart, J., Nunes, D., Rudrapatna, U., Fieremans, E., Jones, D.K., Novikov, D.S., Shemesh, N.. Noninvasive quantification of axon radii using diffusion MRI. eLife 2020;9.

Vincent, M., Palombo, M., Valette, J.. Revisiting double diffusion encoding mrs in the mouse brain at 11.7 t: Which microstructural features are we sensitive to? NeuroImage 2020;207:116399.

Westin, C.F., Knutsson, H., Pasternak, O., Szczepankiewicz, F., Özarslan, E., van Westen, D., Mattisson, C., Bogren, M., O’donnell, L.J., Kubicki, M., et al. Q-space trajectory imaging for multidimensional diffusion MRI of the human brain. Neuroimage 2016;135:345–362.

Wiegell, M.R., Larsson, H.B., Wedeen, V.J.. Fiber crossing in human brain depicted with diffusion tensor MR imaging. Radiology 2000;217(3):897–903.

Yang, D.M., Huettner, J.E., Bretthorst, G.L., Neil, J.J., Garbow, J.R., Ackerman, J.J.. Intracellular water preexchange lifetime in neurons and astrocytes. Magnetic resonance in medicine 2018a;79(3):1616–1627.

Yang, G., Tian, Q., Leuze, C., Wintermark, M., McNab, J.A.. Double diffusion encoding MRI for the clinic. Magnetic resonance in medicine 2018b;80(2):507–520.

Zhang, H., Schneider, T., Wheeler-Kingshott, C., Alexander, D.. NODDI: practical in vivo neurite orientation dispersion and density imaging of the human brain. Neuroimage 2012;61(4):1000–1016.

Zhang, Z., Bassam, B., Thomas, A.G., Williams, M., Liu, J., Nance, E., Rojas, C., Slusher, B.S., Kannan, S.. Maternal inflammation leads to impaired glutamate homeostasis and up-regulation of glutamate carboxypeptidase ii in activated microglia in the fetal/newborn rabbit brain. Neurobiology of disease 2016;94:116–128.

Zhao, C., Teng, E.M., Summers, R.G., Ming, G.l., Gage, F.H.. Distinct morphological stages of dentate granule neuron maturation in the adult mouse hippocampus. Journal of Neuroscience 2006;26(1):3–11.

